# Red seaweed supplementation suppresses methanogenesis in the rumen, revealing potentially advantageous traits among hydrogenotrophic bacteria

**DOI:** 10.1101/2024.06.07.597961

**Authors:** Pengfan Zhang, Breanna Roque, Pedro Romero, Nicole Shapiro, Emiley Eloe-Fadrosh, Ermias Kebreab, Spencer Diamond, Matthias Hess

## Abstract

Macroalgae belonging to the genus *Asparagopsis* have been shown to reduce the production of methane (CH_4_) during rumen fermentation, while increasing feed efficiency when added to the feed of cattle. However, little is known about how the microbial community in the rumen responds to *Asparagopsis* supplementation, and how changes in the rumen microbiome may contribute to shifts in rumen function and ultimately the host’s phenotype. In this study, we generated and analyzed metagenomic and metatranscriptomic data from the microbiome associated with rumen fluid collected from two cohorts of lactating dairy cows, one fed a diet supplemented with *Asparagopsis armata* (treatment) and another fed the same diet without *A. armata* supplementation (control). The reduction of CH_4_ emission from animals that received *A. armata* was coupled to a qualitative decrease in relative archaeal abundance and a significant reduction in the transcription of methanogenesis pathways. Additionally, a significant decrease in the transcription of genes for complex carbon catabolism and a re-organization of the expression profile of carbon catabolic genes at the species level was observed in treated animals. Increased H_2_ production, a consequence of methanogenesis suppression, was coupled to a significant increase in the transcription of hydrogenases that mediate hydrogenotrophic metabolism in the treatment group. Analysis of metatranscriptome data identified a single uncultured hydrogenotrophic bacterial species (a *Duodenibacillus sp.*) as the dominant driver of this transcriptional change. Comparative genomic analysis between the *Duodenibacillus sp.* and other hydrogenotrophic rumen organisms revealed metabolic traits in the *Duodenibacillus* that may provide a competitive advantage in H_2_ scavenging. These findings provide an initial understanding of how rumen microbiota respond to a promising CH_4_ reducing feed additive, and may serve as a model to understand alternative stable rumen microbiome states that produce less methane and increase animal productivity.

## Introduction

Ruminant livestock are a major source of methane (CH_4_), and it has been estimated that they are responsible for nearly 30% of anthropogenic CH_4_ emissions both in the United States and worldwide^1^. Methane, a potent greenhouse gas, exhibits a global warming potential 28 times greater than that of carbon dioxide (CO_2_) over a 100-year period^2^. As the demand for ruminant-sourced products is expected to rise in the coming years, there is an urgent need to develop effective and sustainable approaches to reduce CH_4_ emissions from this sector.

Methane production in ruminant livestock is a byproduct of the anaerobic fermentation of their feed, wherein plant fibers are broken down through a complex synergy among a diverse consortium of microorganisms including bacteria, fungi, and protozoa^3–7^. The rumen microbiome is crucial for the animal’s health and survival due to its key role in generating volatile fatty acids (VFAs), which supply up to 70% of the animal’s energy requirements^8^. Alongside VFAs, this fermentation process also yields substantial amounts of hydrogen (H_2_) and carbon dioxide (CO_2_)^5^. In the terminal stages of rumen fermentation, excess reducing equivalents, such as H_2_, are predominantly eliminated by archaeal methanogens, with a minor contribution from VFA-producing hydrogenotrophic nitrate- and sulfate-reducing bacteria^9^. Despite their relatively lower abundance than bacteria, methanogens serve as the most important electron sink, propelling the fermentation process forward^7,9^. Additionally, by regulating the rumen’s H_2_ partial pressure, methanogens influence the overall pH of the rumen, which affects some of the major fibrolytic rumen bacteria^10^. Consequently the hydrolysis of biomass is likely impacted by methanogen activity, and a notable correlation exists between the abundance of methanogens and cellulolytic microorganisms^11^. Despite their importance as electron utilizers and their broader influences on the rumen microbiome, suppression of methanogens does not always negatively impact rumen functionality^12^. While the various contributions of methanogens could potentially be filled by other naturally occurring rumen microorganisms^13,14^, there is limited data on alternative rumen microbiome states where methanogens are inhibited. This lack of information represents a knowledge gap in rumen microbiology, as the inhibition of methanogenesis could free reducing equivalents for the production of VFAs^15–19^, which in turn would be available for utilization by the host animal, thereby increasing the ruminant’s feed efficiency and productivity^15,18^.

Numerous strategies have been explored to curb CH_4_ emissions from ruminants, including the introduction of biological and synthetic feed additives^15,18,20–24^, dietary adjustments^25–27^, and selective breeding^28,29^. Although selective breeding over several generations has shown to be effective, there is a limitation on the extent it can yield further improvements in the short time frame necessary for climate change mitigation^30^. Conversely, feed additives and dietary modifications offer more immediate avenues for intervention, yet some of them adversely affect VFA production or overall animal productivity^31^. Members of the genus *Asparagopsis,* a red seaweed that produces and stores various halogenated CH_4_ analogues^32,33^, has emerged as a promising biological solution for significantly reducing CH_4_ emissions from ruminants^18,22,31,34^.

Previous work has shown that *Asparagopsis sp.* can cut methane emissions by up to 99% during *in vitro* rumen fermentation without negatively impacting VFA profiles or feed digestibility^31,35^. *In vivo* studies further validate these findings, showing CH_4_ reductions of up to 67% in dairy cows^18^ and up to 98% in beef steers^36^. Notably, the methane-lowering effect of *Asparagopsis* persisted over a prolonged animal trial of 21 weeks and enhanced feed efficiency^37^. This suggests a potential metabolic shift within the rumen microbiome that allowed animals to generate energy precursors without the accompanying methane production.

It has been hypothesized that *Asparagopsis sp.* exert their anti-methanogenic properties likely through the activity of bromoform, or a bromoform derivative, and its capability to bind and inhibit coenzyme F430, which is an essential cofactor in methanogenesis^38,39^. However, the high potency of *Asparagopsis sp.* may stem from the synergistic action of multiple halogenated CH_4_ analogues, targeting various reactions within the methanogenesis pathway. Despite these initial insights, the broader impacts of *Asparagopsis* supplementation on the rumen microbiome, including changes in microbial composition, the regulation of methanogenic pathways, and the broader transcriptional dynamics of the rumen microbiome under methanogenesis suppression, remain poorly understood.

To address this gap, we performed genome-resolved metagenomic and metatranscriptomic analyses of the microbiome that is associated with rumen fluid from dairy cows whose feed were supplemented with *Asparagopsis armata* over 14 days^18^. Our findings revealed qualitative shifts in the rumen microbial community, notably a decrease in methanogenic archaea in animals that received *A. armata* supplemented feed. Furthermore, metatranscriptomic data indicated significant transcriptional reprogramming within the microbiomes of these animals, including: i) a widespread suppression of genes known to be involved in methanogenesis, ii) an upregulation of hydrogenotrophic pathways, and iii) a decrease in the expression of genes associated with complex carbohydrate degradation. These data support the capability of *A. armata*, and most likely of *Asparagopsis sp*. in general, to suppress genes across at least two major methanogenesis pathways. These findings provide foundational and new insights into the mechanisms by which methanogenesis suppression may reorganize rumen microbiome composition and functions towards states that might be more beneficial for the environment and the productivity of the animal. An enhanced understanding of these functional shifts at the genome and gene level will be essential to identify organisms and pathways that drive these functional shifts and to enable the development of next generation rumen modulation approaches.

## Results

### An integrated rumen microbiome database enhances species profiling

Rumen fluid was collected from 8 lactating holstein cows following 14 days of feed supplementation with (Treatment; n = 4 cows) and without (Control; n = 4 cows) *Asparagopsis armata* at an inclusion rate of 1%^18^ (**Figure S1**). Cows treated with *A. armata* exhibited a decrease in methane production of up to 60%, a 367% increase in hydrogen production, and an increase in feed efficiency of up to 74% (**Table S1**). To recover microbial genomes specifically found in the rumen of the animals that were sampled during this study, we extracted microbial DNA from the rumen of one cow from each treatment group and generated ∼ 30Gb of shotgun sequence data per sample for subsequent metagenomic analysis (**Table S2**). Assembly and binning of metagenomic reads from these samples resulted in the recovery of 72 draft genome bins at the species-level (ANI ≥ 95 %) each with estimated completeness ≥ 60% and contamination ≤ 10%. We also generated a total of 400 Gbp (∼ 50 Gb / sample) of metatranscriptome data from all 8 animals in the study (**Table S2**). To increase the probability that metagenomic and metatranscriptomic reads could be assigned to microbial genomes we combined the 72 draft genomes recovered in this study with rumen associated isolate genomes and rumen metagenome-assembled genomes (RuMAGs) from three public datasets^40–42^ to create a comprehensive cohort specific rumen genome database. Initially, this integrated genome database contained 547 genomes from microbial isolates and 10,591 RuMAGs. After these genomes and RuMAGs were de-replicated at the species level (ANI ≥ 95 %) our database contained genomes from 3,119 bacterial and 61 archaeal species across 27 phyla, with 35 species exclusively recovered from this study (**Figure 1, Figure S2, and Table S3**). Hereafter we refer to the 3,180 species-level representative genomes that encompass both isolate assembled genomes and RuMAGs as genomes. Our genome database yielded significantly higher mapping rates for metagenomic samples relative to any of the individual rumen-specific databases alone (**Table S4**). Furthermore, we compared the efficiency with which we could detect microbial species in our samples using a widely used microbiome profiler, MetaPhlAn4^43^, relative to our genome database. We were able to reliably detect more than two times as many species (2,013 vs. 828 species) using our database relative to MetaPhlAn4. This both highlights the importance of comprehensive environment specific genome databases and allowed for significantly improved metagenomic resolution and analysis in this study.

**Figure 1:**
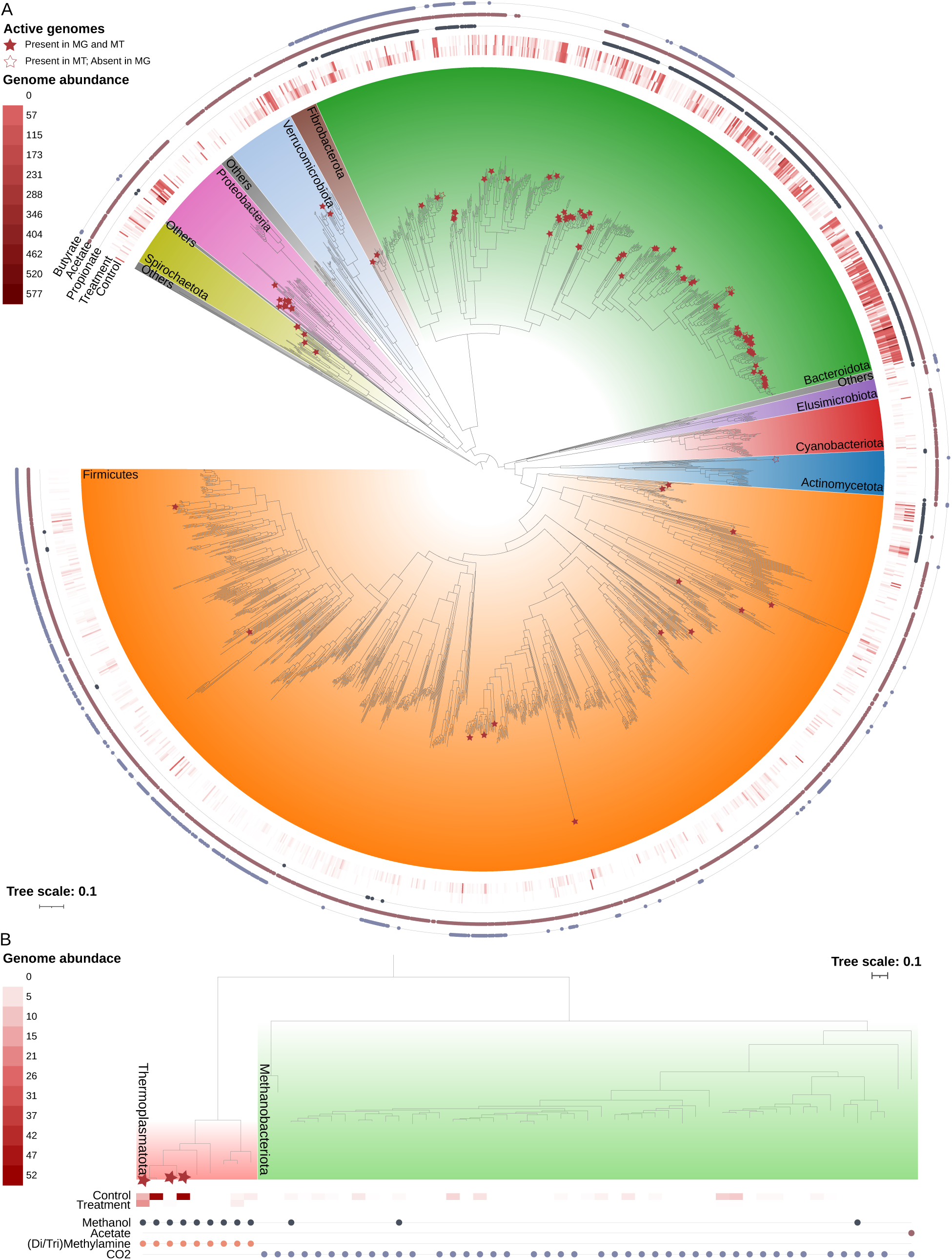
Phylogenetic tree of detected rumen bacteria and archaea. **A**) Phylogenetic tree of 1,949 bacterial genomes detected either in metagenomic or metatranscriptomic samples of this study (n = 2,013 detected in total). Tree was constructed using a concatenated set of bacterial specific marker genes, and rooted at the midpoint. Some species detected in metagenomic or metatranscriptomic samples were omitted from the tree due to insufficient numbers of marker genes. Colored ranges over tree branches indicate phylum level taxonomy. Phyla that had fewer than 20 species were consolidated in “Others”. Solid stars indicate genomes for which were detected in metagenomic samples and ≥ 20% of genes were detected in metatranscriptome samples (n = 99 genomes). Non-solid stars indicate genomes that recruited reads from metatranscriptome samples, but were not detected in metagenome samples (n = 11 genomes). The heatmap shows normalized genome abundance within the metagenome samples as indicated in the inner two rings (for Control and Treatment samples). The outer three rings indicate the presence of VFA (e.g. propionate, acetate and butyrate) biosynthetic pathways in genomes by the presence of colored circles. **B**) Phylogenetic tree of all 58 archeal genomes from the methanogenic phyla *Thermoplasmatota* and *Methanobacteriota* in the rumen specific database created for this study at species level resolution. Tree was constructed using a concatenated set of archaeal specific marker genes, and rooted at the midpoint. Solid stars indicate genomes for which ≥ 20% of genes were detected in metatranscriptome samples (n = 3 genomes). The heatmap shows normalized genome abundance within the metagenomes as indicated in the inner two rings (for Control and Treatment samples). Colored points below the heatmap indicate the presence in genomes of methanogenic metabolic pathways that can utilize different starting substrates. For phylogenetic trees showing all genomes in the rumen specific database created for this study see Supplementary Figure S2.

### Microbial diversity and VFA biosynthetic potential of rumen metagenomes

Metagenomic read mapping against our cohort-specific genome database, revealed a total of 2,013 species-level genomes across the two animals that were profiled, with 1,857 and 1,411 species detected in the control and treatment animal, respectively. Qualitatively, we observed that the 603 species of *Bacteroidota* detected were in relatively high abundance (each species accounting on average for 0.1±0.3% of the observed relative abundance). Additionally, some selected species within the *Pseudomonadota*, *Verrucomicrobia*, and *Fibrobacteria* were also highly abundant (**Figure 1A**) Alternatively, *Firmicutes* were represented by a large taxonomic diversity (1,055 species) but each *Firmicute* species contributed considerably less relative abundance to communities on average (0.01±0.05% average relative abundance per species).

To evaluate the potential of the bacterial rumen population for VFA production, we systematically profiled genomes to look for genes involved in VFA biosynthesis (See Materials and Methods). The potential for acetate biosynthesis was widespread, with the phosphotransacetylase/acetate kinase (PTA/AK) pathway being the dominant route (**Figure 1A and Figure S2A**). Alternative routes for acetate biosynthesis via the Acetyl-CoA (generating ADP) and Butyryl-CoA pathways were also detected specifically in *Bacteroidota*. Acetate biosynthesis through the Acetyl-CoA synthetase (ACS) route generating AMP was rare in genomes, and the potential for propionate and butyrate biosynthesis was found to be clade-specific (**Figure 1A and Figure S2A**). Propionate biosynthetic pathways were primarily found among the *Bacteroidota*, with the succinate-mediated pathway being the primary route. Alternative pathways for propionate biosynthesis were only detected in a limited number of genomes and pathways for butyrate biosynthesis were primarily detected in subgroups of *Firmicutes* and *Bacteroidota*. Unlike genomes encoding pathways for acetate and propionate production, we found that genomes encoding butyrate production pathways frequently encoded multiple routes. Specifically, we noted the co-existence of lysine- and pyruvate-mediated butyrate pathways in *Bacteroidia* genomes. Taken together, the observed relative abundance of bacterial species and the VFA production potential suggest that *Bacteroidota* species likely play an important role in VFA production in these samples, particularly for butyrate biosynthesis.

### Metatranscriptomic profiling captures relevant bacterial and archaeal species

Transcriptome profiling of all 8 rumen fluid samples collected during this study identified transcripts from 1,865 microbial species across all samples, including 1,784 and 1,254 species in the control and treatment rumen microbiome respectively. However, most of the detected genomes recruited only a small number of transcripts suggesting low transcriptional activity, with the majority of transcripts (73.2±10.8%) mapping to a core set of 110 bacterial and 3 archaeal species that were considered transcriptionally active (≥ 20% of their genes transcribed in at least one sample). These 113 species broadly encompass organisms detected at high relative abundance in our metagenomic samples and spanned 7 bacterial and 1 archaeal phyla (**Figure 1 and Figure S2**). In the control group, two species belonging to the *Enterobacterales* and *Bacteroidales* were highly transcriptionally active, contributing 18.4±16.5% and 11.1±3.2% of all mapped metatranscriptome reads respectively. Alternatively in treatment samples, the 10 most abundant genomes recruited on average a similar fraction (3.0±1.1%) of metatranscriptome reads. Overall, a comparison between the rumen metatranscriptome profiles of treated and untreated animals found a statistically significant separation based on treatment (p-val = 0.034; PERMANOVA) between these sample sets (**Figure S3**).

### *Asparagopsis armata* treatment decreases archaeal abundance and transcription

Within the metagenomes, *Methanobrevibacter sp.* from the *Methanobacteriota* and *Methanomethylophilus sp.* from the *Thermoplasmatota* were the most abundant archaeal methanogens detected (**Figure 1B**). Functional profiling of methanogenesis pathways across these two taxa confirmed that *Methanobrevibacter sp.* genomes encoded the genes to perform hydrogenotrophic methanogenesis while *Methanomethylophilus sp.* genomes encoded the genes necessary for methylotrophic methanogenesis, which involves methanol and methylated amine reduction (**Figure 1B**). Our metagenomic data also showed that *Methanomethylophilus sp.* were the dominant members within the archaeal methanogen populations, accounting for 66.7% and 82.2% of the archaeal relative abundance in the control and treatment metagenomes respectively. Previous 16S rRNA amplicon based rumen surveys^44^ suggested that hydrogenotrophic methanogens of *Methanobrevibacter sp.* dominate methanogenic rumen populations, accounting for ∼74% of the 16S rRNA gene amplicons. This discrepancy in archeal species abundance between studies may be due to the fact that *Methanobrevibacter sp.* possess 2-3 copies of the 16S rRNA gene whereas *Methanomethylophilus sp.* possess 1 copy^45^, leading to an overestimation of *Methanobrevibacter sp.* using 16S rRNA profiling methods. Alternatively, shotgun metagenomics is agnostic to uneven copy numbers across genomes when abundances are quantified using metagenome assembled genomes^46^.

Metagenomic profiling revealed that treatment with *A. armata* resulted in a reorganization of the rumen microbiome including a qualitative increase of *Bacteroidota* and decrease of *Pseudomonadota* within the treated animal relative to the control animal (**Figure 2A**). Moreover there was a reorganization of the most abundant species in the control and treatment rumen microbiomes (**Figure S4**). There was a notable qualitative effect on archaeal populations, which decreased in abundance from 1.76% (control) to 0.02% (treatment) (**Figure 2B**). In particular *Methanobacteriaceae sp.* were relegated almost undetectable, and we observed that the relative abundance of almost all individually profiled archaeal genomes fell to near zero with exception of two *Methanomethylophilaceae sp.* genomes (**Figure 2C**).

**Figure 2:**
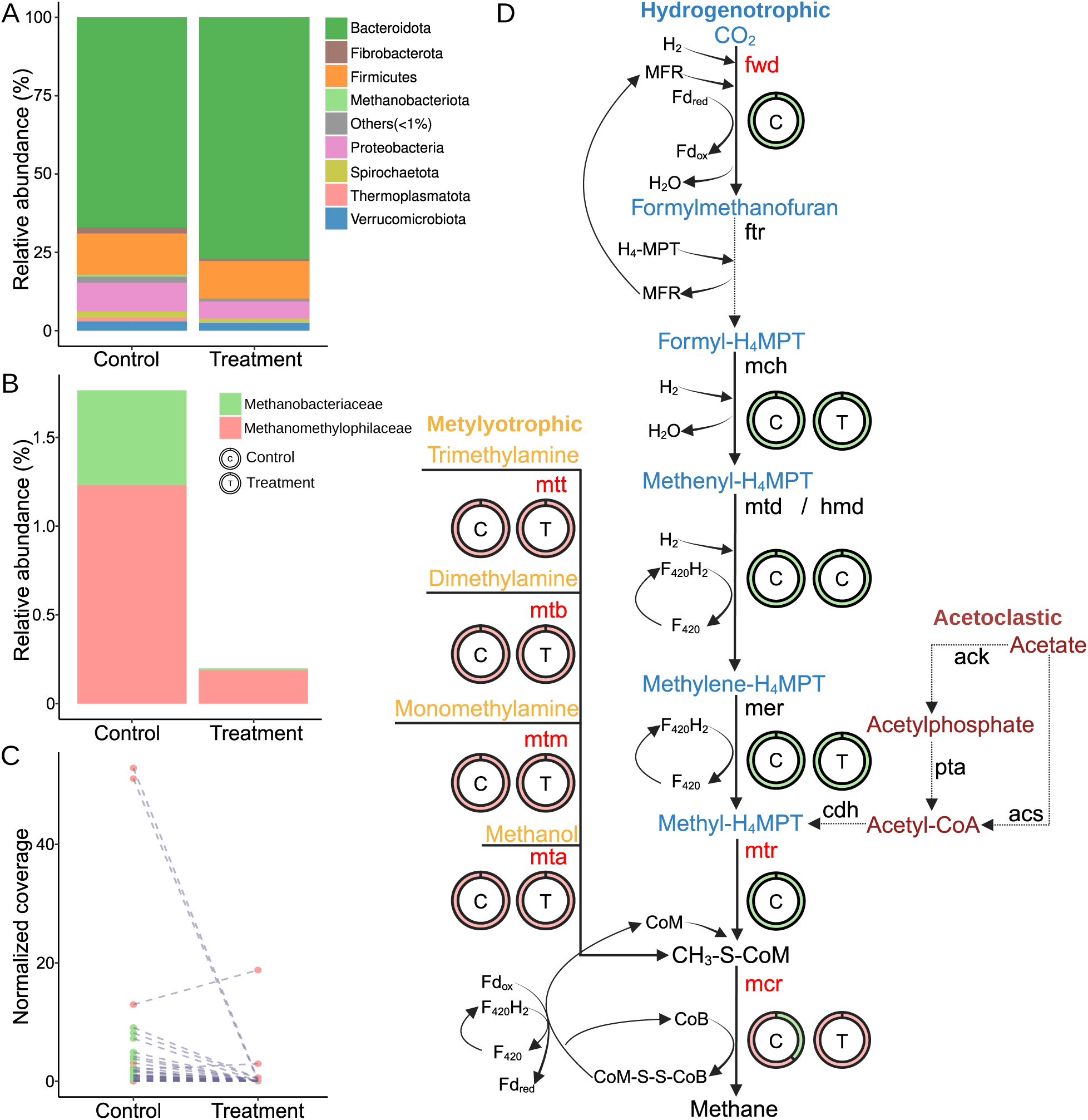
*A. armata* supplementation impacts on microbial abundance and methanogenic pathways. **A)** Microbial relative abundance at the phylum level from metagenomic samples. Bacterial phyla whose relative abundance is < 1% are labeled as ‘Others’. **B)** Archaeal relative abundance at the family level from metagenomic samples. **C)** Coverage of individual archaeal genomes between treated and control metagenomic samples. Lines connect the same genome detected in control and treatment samples. **D)** Gene expression changes across 3 profiled methanogenic pathways, with the names of metabolic intermediates colored based on the type of methanogenic pathway (methylotrophic - yellow, hydrogenotrophic - blue, acetoclastic - red). Solid and dotted black lines indicate transcripts for an enzyme were or were not detected in samples, respectively. Enzyme names depicted in red indicate that at least one enzyme subunit was significantly down-regulated in treated samples (and no subunit was up-regulated). Enzyme names in black showed no statistically significant difference between samples. Circles show the relative contribution of archeal families to the transcriptome reads of each enzyme in both control (C) and treated (T) samples.

The qualitative reduction of archaeal populations, in samples treated with *A. armata*, paralleled a broad and significant decrease of transcripts associated with methanogenesis pathways (**Figure 2D**). This included significant suppression of transcription of the formylmethanofuran dehydrogenase subunit A (fwdA; p-val = 0.046; log_2_FC = -5.8) and the methyltransferase subunits A and H (mtrAH; p-val = 0.046/0.039; log_2_FC = -5.3/-5.6) and transcripts for the methyl coenzyme M reductase enzyme (mcrABG; p-val = 0.09/0.00001/0.039; log_2_FC = -3.1/-7.9/-5.6). Additionally, while several genes involved in hydrogenotrophic methanogenesis were not detected as significantly down-regulated, all exhibited decreases in transcript abundance in treated samples (**Table S5**). Consistent with our metagenomic analysis (**Figure 1B and Figure S2B**), transcripts for acetoclastic methanogenesis were not detected in our metatranscriptomic data (**Figure 2D**).

### Hydrogenotrophic metabolism is activated in *A. armata* treated animals

Roque, et al. observed that the significant decrease in CH_4_ production in. *A. armata* treated animals was coupled to a significant increase in H_2_ emissions^18^ (**Table S1**). To evaluate how higher H_2_ partial pressures may impact rumen microbiome activity, we specifically assessed the expression of the catalytic subunit proteins for 38 known hydrogenase subgroups^47,48^ between treated and control samples.

Hydrogenase families associated with fermentative H_2_ evolution, specifically FeFe_A1 and FeFe_B, were actively and highly transcribed by rumen bacteria yet showed no significant differences in mean transcript levels between treated and untreated samples (**Figure 3A**). We found that the majority of FeFe_A1 family transcripts originated from only two genomes, a *Succinivibrionaceae* (MGYG000292509) and a *Ruminococcaceae* (MGYG000291593) together contributing 78.3±10.8% and 81.8±2.2% of these transcripts on average in control and treatment samples, respectively (**Figure S5A**). Similarly FeFe_B family transcripts largely originated from a small number of species including a *Treponematales* (RuMAG_Treponema_1) contributing 46.3±10.6% of these reads on average in control samples, and 4 *Bacteroidales* species contributing 78.8±25.3% of these reads on average in treated samples (**Figure S5B**). Overall these results suggest that transcription of H_2_ evolving hydrogenases is dominated by a small number of species in our samples and generally unaffected by treatment with *A. armata*.

**Figure 3:**
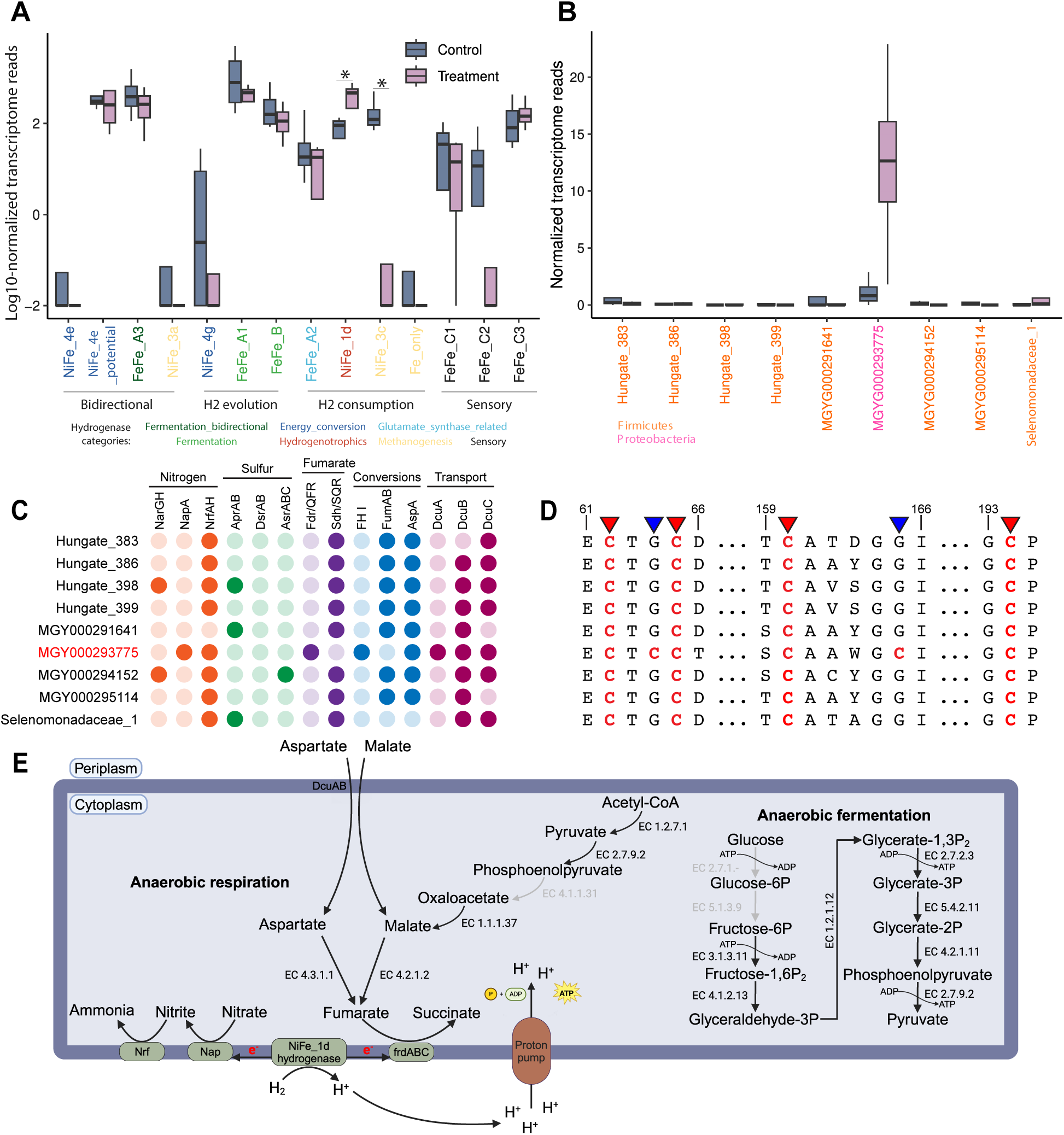
A. armata impacts on hydrogen metabolism and metabolic analysis of hydrogen utilizing species. **A**) Transcriptional changes of 14 hydrogenase families with detectable expression between control and treated samples. The y-axis represents log_10_-normalized transcriptomic reads (per million reads) mapped to the catalytic subunit representatives of each hydrogenase subgroup detected in genomes. Hydrogenase families are colored and organized by the direction of hydrogen flow. Asterisks indicate significant differences in expression between treatment and control (FDR ≤ 0.05; DESeq2). **B**) Normalized counts of transcriptome reads contributing to NiFe-1d expression in 9 species encoding and expressing this hydrogenase family. **C**) Comparative genomic analysis of the 9 NiFe_1d encoding species focused on terminal electron accepting pathways. Potential pathways are organized by nitrogen metabolism (orange), sulfur metabolism (green), fumarate metabolism (purple), fumarate interconversions (blue) and metabolite transport (magenta). The presence of a function in a genome is indicated by a dark circle, while lack of function is indicated by a light circle. NarGH: dissimilatory nitrate reductase; NapA: periplasmic nitrate reductases; NrfAH: ammonia-forming nitrite reductase; AprA: adenylylsulfate reductase; DsrAB: dissimilatory sulfite reductase; AsrABC: anaerobic sulfite reductase; Fdr/QFR: quinol:fumarate reductase; Sdh/SQR: succinate dehydrogenase/succinate-ubiquinone oxioreductase; FH I: fumarate hydratase class I; FumAB: fumarate hydratase; AspA: aspartate-ammonia lyase; DcuABC: C4-dicarboxylate antiporters. **D**) Portion of a multiple sequence alignment of NiFe_1d small subunit proteins from the 9 species in panel C. Proteins are associated with species in the same order as shown in panel C. This alignment depicts the proximal iron-sulfur cluster binding motif in the NiFe_1d small protein subunit. Metal binding cysteines are colored red in the alignment. The 4 cystines conserved across all proteins, which are typically involved in the formation of a 4Fe-4S cluster are highlighted by red triangles. The substitution of 2 cysteines with 2 glycines that differentiate MGYG000293775 from other species, enabling the formation of a 4Fe-3S cluster, are highlighted by blue arrows. Alignment positions are noted above the alignment. Also see supplementary datasets. **E**) Metabolic reconstruction of potential energy generating pathways in MGYG000293775 focused on H_2_ coupled fumarate reduction. Black and gray arrows represent presence or absence of genes encoding reactions, respectively.

We identified two hydrogenase families (i.e., NiFe_3c and NiFe_1d) associated with hydrogen consumption that significantly differed in overall expression between treatment and control samples. Furthermore, we observed that H_2_ consuming hydrogenases contributed a relatively larger fraction of transcripts to the total pool of hydrogenase transcripts in *A. armata* treated samples (control = 18.1±7.3%; treatment = 36.8±5.5%; **Figure S6A**). The NiFe_3c hydrogenase family, which is involved in methanogenesis, was significantly down-regulated (p-val = 0.004; log_2_FC = -4.5) in treated samples (**Figure 3A**). Alternatively, the NiFe_1d hydrogenase family, reported to be involved in hydrogenotrophic metabolism^48^, was significantly up-regulated (p-val = 0.04; log_2_FC = 2.1) in treated samples (**Figure 3A**). Out of 9 genomes expressing NiFe_1d hydrogenases, we observed transcripts were primarily derived from a single *Duodenibacillus sp.* genome (MGYG000293775; control = 37.4±39.8%; treatment = 95.7±4.3%) within the *Pseudomonadota* (**Figure 3B**). Our metagenomic analysis also showed that the relative abundance of MGYG000293775 increased by 58.3-fold in treatment relative to control samples.

### *Duodenibacillus sp.* encodes advantageous energy generating pathways

The increases in relative abundance and gene expression of MGYG000293775, a hitherto uncultured bacteria classified as a *Duodenibacillus sp.*, prompted us to search for metabolic traits that may confer an advantage to this hydrogenotrophic microbe when competing for hydrogen as electron donor. We comparatively evaluated the potential of all 9 transcriptionally active, NiFe_1d encoding, hydrogenotrophic genomes to couple hydrogen oxidation to a number of anaerobic terminal electron acceptors (**Figure 3C**). Among these, 3 genomes encoded genes for dissimilatory sulfur reduction, and 8 encoded genes for dissimilatory nitrogen reduction. However, only 3 genomes, including MGYG000293775, had the potential for complete reduction of nitrate to ammonia encoding both a nitrate reductase (NapA or NarG) and nitrite reductase (NrfA). Distinctively, MGYG000293775 was the only species encoding a periplasmic nitrate reductase (NapA), which is speculated to exhibit increased nitrate affinity^49^, and a quinol:fumarate reductase (FdrA/QFR) capable of utilizing fumarate as a terminal electron acceptor and H_2_ as the corresponding electron donor^50,51^. The succinate-ubiquinone oxidoreductase enzymes (SdhA/SQR) encoded in the other 8 genomes preferentially oxidize succinate to fumarate, and do not typically enable the use of fumarate as a terminal electron acceptor^52–54^. Moreover, we observed that expression of all quinol:fumarate reductase subunits (FdrABC), nitrate reductase (NapA) and nitrite reductase (NrfA) in MGYG000293775 increased in treatment relative to control samples (**Figure S6B**).

Given the important role of fumarate in propionate biosynthesis^55^, free fumarate may be scarce in the rumen environment. In the human gut, where freely available fumarate is also limited, some species support H_2_/fumarate respiration through the import of C4-dicarboxylates (e.g. malate and aspartate) via C4-dicarboxylate membrane transporters and the subsequent conversion of the imported C4-dicarboxylates into fumarate, while simultaneously exporting succinate^56^. Accordingly, we evaluated if dcu genes encoding these C4-dicarboxylate/succinate antiporters were present in the 9 hydrogenotrophic genomes evaluated. We found that MGYG000293775 encoded 8 C4-dicarboxylate antiporter (Dcu) genes, more than all other hydrogenotrophic genomes, including a DcuA variant that was absent from the other 8 genomes (**Figure 3C**). Moreover we found that Dcu transporters in the MGYG000293775 genome were co-located with genes that may support H_2_/fumarate respiration by mediating the conversion of C4-dicarboxylates to fumarate (**Table S6**). This included the co-location of *dcu*A with an aspartate-ammonia lyase (*aspA*), the proximal co-location of *dcu*B (i.e., within 11 genes) from a fumarate hydratase (*FH I*), collectively enabling the conversion of both L-aspartate and L-malate into fumarate. While the other 8 hydrogenotrophic genomes evaluated encode aspA and fumA, these were not co-located with dcu transporters.

As sequence variation can impact the activity and oxygen tolerance of NiFe_1d hydrogenases^57,58^, we evaluated the sequence similarity of these hydrogenases across all 9 NiFe_1d encoding genomes (**Supplementary Datasets**). We found that the essential L1 (xxRI**C**GV**C**TxxH) and L2 (SFDP**C**LA**C**xxH) motifs in the NiFe_1d large subunit were completely conserved across sequences from all of the 9 NiFe_1d encoding genomes in our analysis. However the NiFe_1d small subunit protein of MGYG000293775 was the only protein that contained 6 cysteines in the proximal iron-sulfur cluster motif (E**C**Tx**CC**…x**C**AxxG**C**…G**C**P) compared to 4 cysteines (E**C**Tx**C**x…x**C**AxxGx…G**C**P) observed in the other small subunit proteins (**Figure 3D and Supplementary Datasets**). The presence of 6 cysteines enables the formation of an oxygen tolerant 4Fe-3S cluster, as opposed to the formation of a 4Fe-4S cluster when 4 cystines are present, and may have an impact on the enzymatic reaction rate of the NiFe_1d hydrogenase^47,57^. Taken together, these findings show that MGYG000293775 possesses a multitude of metabolic traits (**Figure 3E**) that may be potentially advantageous, and provide a competitive edge relative to other hydrogenotrophic rumen microbes.

### *A. armata* treatment suppresses expression of carbon catabolic genes

Syntrophic interactions between bacteria and methanogens have been reported^59^, and in culture H_2_ partial pressure has been shown to impact rumen bacterial cellulolytic activity^60^. Given the strong impact of *A. armata* treatment on the abundance and activity of rumen methanogens (**Figure 2B-D**), we evaluated the expression of genes involved in the degradation of complex plant carbohydrates. Differential gene expression profiling revealed 25 Carbohydrate Active Enzyme (CAZymes) families with significant expression differences between treated and untreated animals (**Figure 4A and Table S7**). Of these 25 CAZyme families, all but one, CAZyme family GH11, were significantly down-regulated in the microbiomes of ruminants that received *A. armata* (**Figure 4A**). The CAZyme family most strongly down-regulated was Glycoside Hydrolase Family 48 (FDR = 2.4e^-4^; Log_2_FC = -5.35), which includes chitinase, cellobiohydrolase and cellulase enzymes, known to significantly contribute to fiber degradation in the rumen^6,61,62^. Numerous other CAZyme families targeting plant polymers including hemicelluloses (GH26 and GH130) and pectins (PL9, PL11, and GH106) were also transcriptionally downregulated. Xylanases, belonging to CAZyme family GH11, were the sole GH family upregulated in the treatment group. GH11 transcripts were primarily derived from *Bacteroidota* and *Fibrobacterota* in control samples, and this shifted towards more dominant expression from *Bacteroidota* in treated samples (**Figure S7**). Members of the GH11 family from *Fibrobacter succinogenes* were enriched in the metaproteome of the fiber adherent rumen microbiome^6^ and expression levels of this family have been reported to be higher in feed efficient animals^63^.

**Figure 4:**
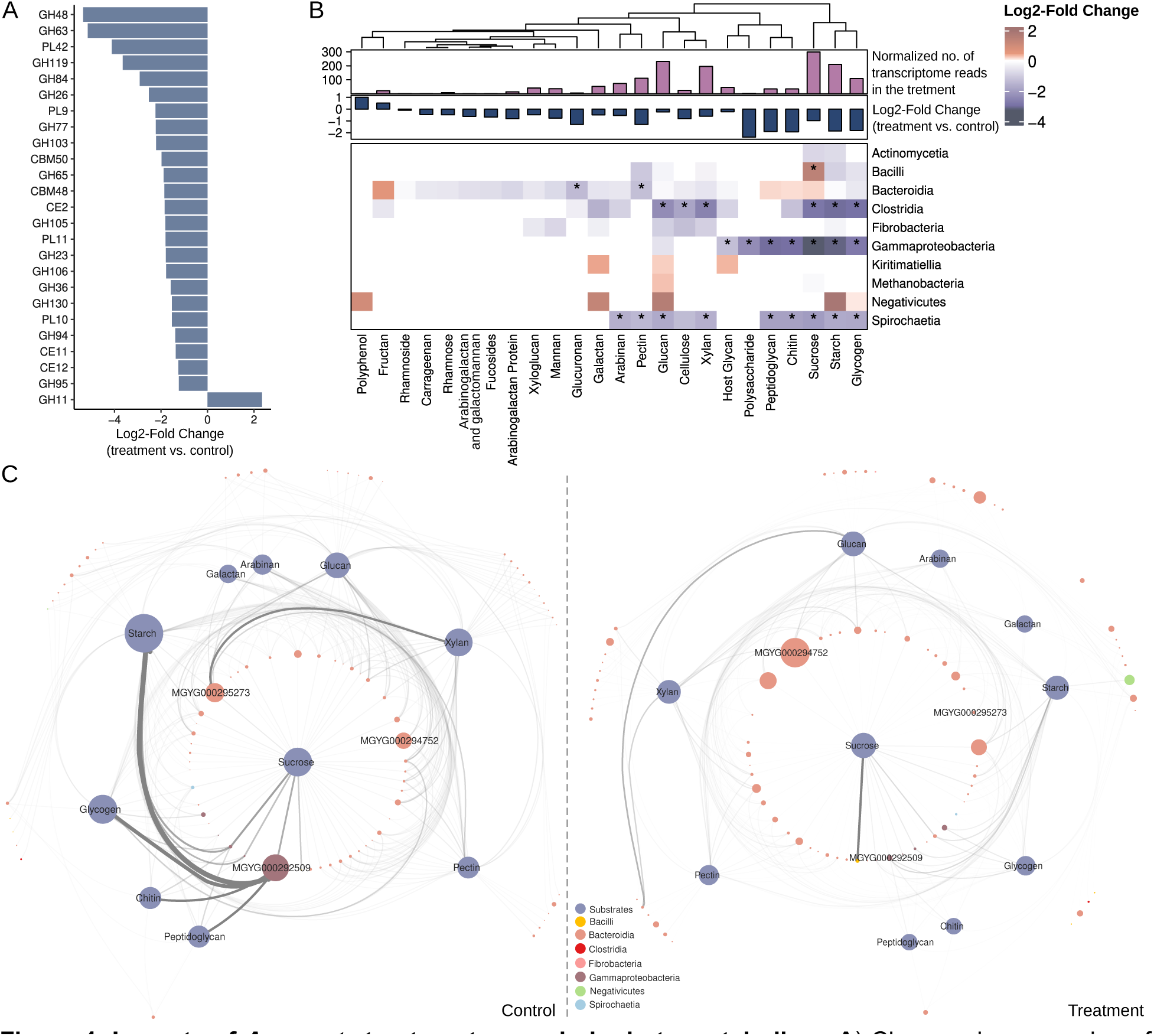
Impacts of *A. armata* treatment on carbohydrate metabolism. **A**) Changes in expression of 25 CAZyme families with significant differential expression between control and treated samples. The x-axis shows log2-fold change from DESeq2 (FDR ≤ 0.05). The y-axis shows individual CAZyme families. Glycosyl Transferases (GT) were omitted from the analysis. **B**) Changes in expression of CAZYmes grouped by order-level microbial taxa and potential substrate. If a CAZyme family acted on more than one substrate it was counted for both. The heatmap represents log2(treatment/control) change of normalized transcripts assigned to CAZymes degrading a specific substrate in each class. The bar plots above the heatmap show the total normalized number of transcripts (per million reads) assigned to CAZymes targeting each substrate (upper, purple), and log2(treatment/control) change of normalized transcripts assigned to specific substrates (lower, blue). **C**) The top 250 genomes and top 10 substrates, assessed by total CAZyme transcription in either control (left panel) or treatment (right panel) samples. Genomes (non-blue nodes) are connected to substrates (blue-nodes) if they express CAZymes active on that substrate. The width of network edges denotes the expression level of CAZymes targeting a substrate. The size of the carbohydrate nodes represent the total normalized number of reads assigned to the cazymes targeting the specific substrate.The size of the genome nodes represents the genome abundance in either the control or treatment sample measured by metagenomics.

Next, we evaluated the relative contribution of the rumen microbiome, at class-level resolution, to the expression of CAZymes and their potential substrates (**Figure 4B**). Transcripts from CAZymes with substrate specificity towards common plant biomass carbohydrates, including sucrose, starch, xylan, glucan, and pectin, were derived from microbes across numerous taxonomic classes, highlighting the overall importance of these CAZymes for rumen biomass degradation. However, the number of CAZyme transcripts declined across almost all substrate categories and taxonomic classes in the rumen microbiomes of animals fed an *A. armata* supplemented diet (**Figure 4B**). Members of the *Bacteroidia*, *Clostridia*, *Gammaproteobacteria*, and *Spirochaetia* exhibited transcriptional activity across the largest set (≥7) of potential substrates. However CAZyme transcripts primarily from *Clostridia*, *Gammaproteobacteria*, and *Spirochaetia* were significantly down-regulated across most potential substrates in the rumen microbiomes subjected to *A. armata* (**Figure 4B**). Despite the overall decrease in CAZyme expression across taxonomic groups in the presence of A. armata, the relative contribution of different taxonomic classes to substrate-specific CAZyme groups remained consistent (**Figure S8**). Furthermore, CAZyme downregulation was minimal for *Bacteroidia* and not observed for *Fibrobacteria*, a taxonomic group that includes cellulolytic rumen bacteria such as *Fibrobacter succinogenes* S85 (*Fibrobacter succinogenes*).

The consistency of the relative contributions of class-level microbial lineages to CAZyme expression between treatment and control samples (**Figure S8**) prompted us to investigate if there was a reorganization of CAZyme gene expression at the species level. In control samples, two species from *Succinivibrionaceae* (MGYG000292509) and *Prevotella* (MGYG000295273) respectively displayed the highest CAZyme expression (**Figure 4C**). These organisms were also the two most abundant in the control sample we assessed with metagenomics. MGYG000292509 expressed CAZymes linked to starch, glycogen, peptidoglycan and chitin degradation, while MGYG000295273 expressed CAZymes primarily linked to xylan degradation (**Figure 4C**). In samples treated with *A. armata*, we observed significantly decreased CAZyme transcription from both of these species. Moreover, a variety of genomes from other species became the dominant contributors to CAZyme gene expression, and we observed a species level reorganization of CAZyme gene expression despite consistent class-level contributions (**Figure 4C and Figure S8**).

### The influence of *A. armata* on VFA biosynthesis

Previous work has shown that supplementation with *Asparagopsis sp.* does not decrease VFA titers *in vitro*^64,65^, and has no negative impact on animal feed conversion efficiency^18^ (**Table S1**). To investigate the effect of *Asparagopsis sp.* supplementation on the biosynthesis of prominent VFAs (e.g. acetate, butyrate and propionate) we profiled the expression of the genes mediating the biosynthesis of these compounds (**Figure 5 and Table S8**).

**Figure 5:**
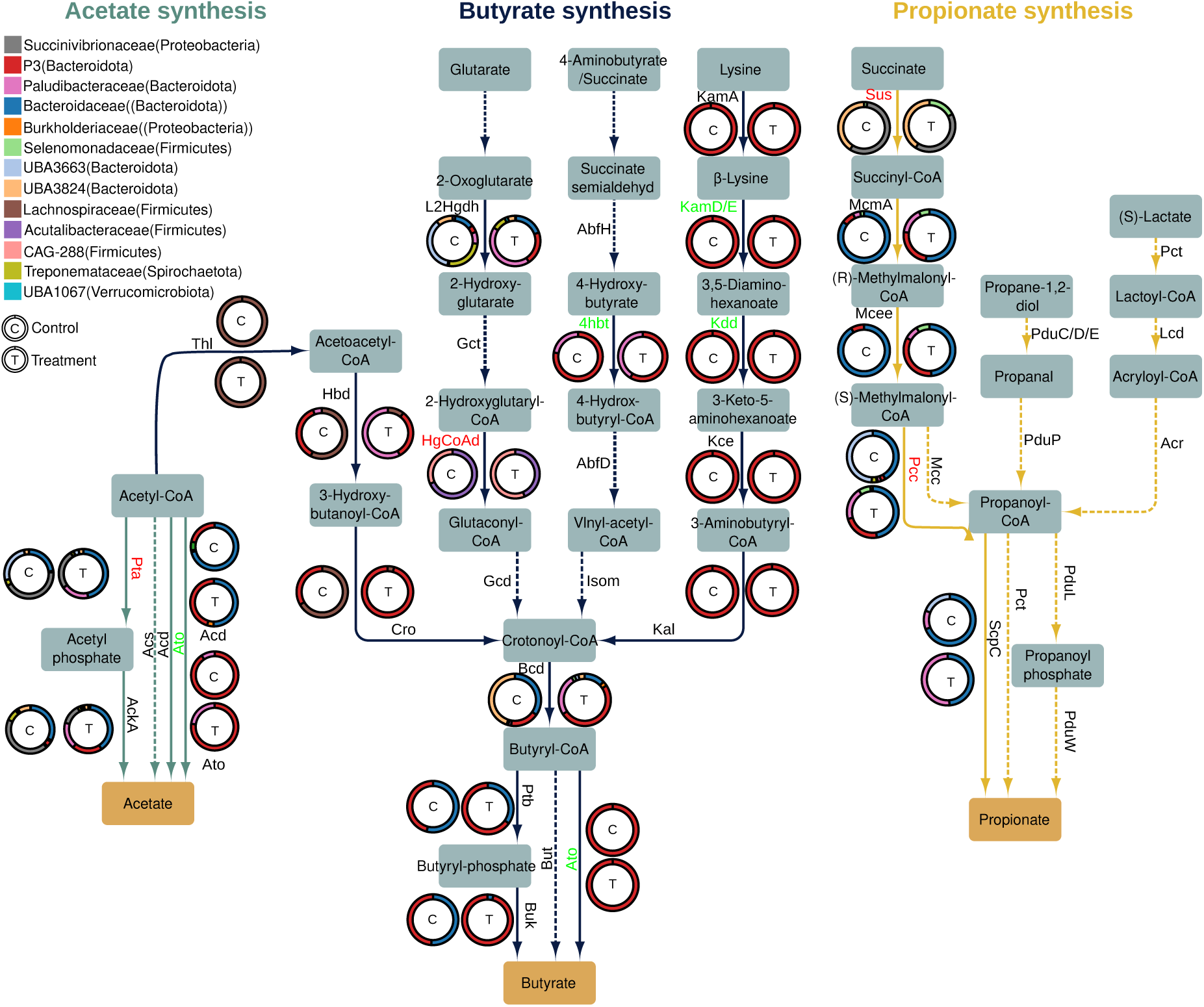
Impact of *A. armata* treatment on transcription of VFA biosynthetic pathways. Pathways and intermediates for acetate, butyrate and propionate biosynthesis are shown. Arrows indicate individual reactions and gene names for protein products mediating each reaction are shown next to arrows. Solid and dashed arrows indicate that a gene product was or was not detected in our metatranscriptomic data, respectively. Arrows are colored based on the presence of reactions as part of acetate (teal), butyrate (blue) and propionate (yellow) biosynthesis. Gene names are depicted in green or red to indicate significant up-regulation or down-regulation (FDR ≤ 0.05) in the treatment condition, respectively. For reactions with multi-subunit enzymes to be considered up or down-regulated, at least one subunit had to be differentially expressed and no other subunit could have the opposite direction of expression. The rings present next to each gene product give the fraction of transcriptomic reads associated with genomes of different taxonomic families that are assigned to each enzyme in control (C) and treatment (T) samples. For taxonomic profiles the number of reads from all gene subunits were aggregated. Also see Table S8.

In the biosynthesis of acetate we observed transcriptional changes in two biosynthetic routes, the *pta/ackA* (phosphate acetyl-transferase/acetate kinase) mediated route where acetyl phosphate is an intermediate, and the Acetyl-CoA to acetate route mediated by *ato* (butyryl-CoA:acetoacetate CoA transferase). The expression of *pta* was primarily associated with genomes of the *Bacteroidaceae* and *Succinivibrionaceae* taxonomic families, and *pta* transcription was significantly down-regulated (p-val = 2e^-6^; Log_2_FC = - 2.0) in the rumen microbiome subjected to *A. armata* (**Figure 5**). Furthermore, we observed that in treated samples *pta* transcripts from the *UBA3663* and *Paludibacteraceae* families contributed to a lower and higher fraction of overall pta transcripts, respectively. The expression of *ato* was primarily associated with genomes of the *Paludibacteraceae* and *P3* taxonomic families, and *ato* transcription was Psignificantly up-regulated (p-val = 0.002; Log_2_FC = 2.5) in the rumen microbiome subjected to *A. armata* (**Figure 5**). No major changes to the taxonomic distribution of *ato* transcripts was observed.

Transcripts for genes that mediate butyrate biosynthesis from 4 independent precursors, acetoacetyl-CoA, glutarate, 4-aminobutyrate/succinate, and lysine were detected in our samples (**Figure 5**). In the lysine derived butyrate pathway we observed a significant increase in the transcription of genes for three steps, *kamDE* (β-lysine-5,6-aminomutase; p-val = 0.0036; log_2_FC = 1.7), *kdd* (3,5-diaminohexanoate dehydrogenase; p-val = 0.0036; log_2_FC = 1.99), and *ato* (p-val = 0.0003; log_2_FC = 2.4) in treated samples (**Figure 5**). Due to *ato* functioning in both acetate biosynthesis and the lysine-dependent butyrate biosynthesis pathways, respectively^66,67^, we evaluated *ato* transcripts associated with the lysine-dependent butyrate pathway by only including those from organisms that expressed full sets of genes for this pathway (i.e. *kamA*, *kamDE*, *kdd*, *kce*, *kal*, *bcd*, and *ato*).

Expression of genes associated with propionate biosynthesis were exclusively detected for the succinate-mediated route (**Figure 5**). The expression of both *sus* (Succinyl-CoA synthetase; p-val = 0.001; log_2_FC = -2.6) and *pcc* (propionyl-CoA carboxylase; p-val = 0.039; log_2_FC = -0.9) were significantly decreased in microbiomes of the treatment group. Taken together, we found that genes mediating VFA biosynthesis were primarily expressed from genomes in the phylum *Bacteroidota*, and that while most detectable genes for VFA biosynthesis were not differentially expressed, a limited set of potentially important transcripts were differentially expressed under methanogenesis suppression.

## Discussion

Despite the effort towards developing advanced strategies for curbing the production and emission of methane (CH_4_) from cattle^7,68^, these interventions often only lead to a partial reduction of CH_4_ emissions^7,69^ and fail to reroute methanogenic substrates towards alternative metabolites^31^, which could enhance the productivity of the ruminant animal^8^. Developing approaches for targeted and lasting modulation of the rumen microbiome, in a manner that both diminishes CH_4_ output and bolsters animal production efficiency, necessitates a detailed microbiome understanding of this phylogenetically complex^44^ and metabolically interconnected environment^70,71^. This includes insights into the complex interactions among its diverse metabolic guilds and their collective response to the suppression or loss of methanogenic activity^72^. To further these goals we performed a species-level metagenomic and metatranscriptomic evaluation of *A. armata* supplemented feeding on the rumen microbiome, a treatment previously shown to potently suppress CH_4_ emissions and increase feed efficiency, which is defined as protein yield per dry matter intake^18^. Despite limited cohort size, our research revealed: (i) a pronounced reduction in both the abundance of methanogens and their transcriptional activity; (ii) a widespread decrease in the transcription of CAZymes; and (iii) the initiation of hydrogenotrophic pathways alongside the identification of species-specific metabolic characteristics that may offer a competitive edge to specific hydrogenotrophic species within the complex rumen ecosystem.

Interrogation of the rumen microbiome at a species level has been bolstered by significant efforts to both cultivated rumen microorganisms^40^ and recover representative sets of rumen genomes for both cultivated and uncultivated species^40,41,62,73^. Despite the availability of several rumen specific genome databases^40–42^, we observed that microbial diversity in our samples remained underrepresented, with read mapping rates ranging from 1% to 18% using these existing databases (**Table S4**). This suggests that rumen microbiomes, even from cattle housed in industrialized countries and subjected to more standardized farming practices, still contain a large and unexplored genomic diversity, with bacterial groups in particular likely remaining undersampled. Despite the limited metagenomic sampling of this work, our newly recovered genomes contributed 32 novel species to our integrated database of 3,119 bacterial and 61 archaeal species. Importantly we observed that the integration of multiple databases increased our ability to map metagenomic reads (**Table S4**) and enabled the interrogation of VFA biosynthetic potential across currently available rumen bacterial species (**Figure 1A and S2A**), which to our knowledge has not previously been done at this scale. An important insight from this analysis is that rumen bacteria utilize numerous pathways for VFA biosynthesis with a taxonomically diverse set of species across the phylum *Bacteriodota* encoding the majority of these pathways (**Figure 1A and S2A**). Furthermore, metatranscriptomic analysis indicated that *Bacteriodota* were also the most active propionate, butyrate and acetate producers in the rumen samples we evaluated (**Figure 5**).

In our study, the dominant methanogenic archaea identified were *Methanomethylophilus* sp. and *Methanobrevibacter* sp. (**Figure 1B**), which aligns with previous work^44^. While traditionally *Methanobrevibacter sp.* have been reported as more prevalent, constituting up to 74% of rumen archaeal populations^44^, our analysis revealed a predominance of *Methanomethylophilus sp.* This deviation may be attributed to a difference in 16S rRNA gene copy number, as *Methanobrevibacter sp.* harbor 2 to 3 copies, versus a single copy in *Methanomethylophilus sp.*^45^. These differences can only be taken into consideration when determining abundances using whole genomes. This observation underscores the importance of using genome-resolved methods, over traditional 16S rRNA amplicon sequencing to accurately quantify methanogenic archaea, given the latter’s susceptibility to gene copy number variations^74^. Utilizing, or constructing *de novo*, appropriate reference genome databases that contain taxonomically comprehensive sets of archaeal genomes associated with the rumen environment is critical. A previous effort failed to detect methylotrophic methanogenic pathways in rumen metagenomes^75^, later shown to be due to the absence of the corresponding genomes from the reference database employed^76^. *Methanomethylophilus sp.* represent a relatively novel and less understood group of methanogens^77^ and their level of abundance in this study suggests that these lesser-studied microbes could play a significant role in rumen methane emissions in some cattle populations. However, our findings, based on metagenomic and metatranscriptomic data (**Figure 2**), indicate that *A. armata* supplementation can effectively suppress both hydrogenotrophic and methylotrophic methanogenesis, concurrently reducing the populations of both *Methanomethylophilus* sp. and *Methanobrevibacter* sp. (**Figure 2B-C**).

The decrease in methanogen abundance and activity (**Figure 2**), paired with a rise in hydrogen production (**Table S1**), aligns with the established theory that methanogens are the most important hydrogen sink in the rumen. However, the broader responses of the rumen microbiome to the loss of this hydrogen sink are poorly characterized. Despite examining the effects of a single additive, our study directly assessed this dynamic, allowing the evaluation of rumen hydrogen metabolism under methanogenesis suppression. While we found no significant changes in fermentative H_2_-evolving or bi-directional hydrogenase expression, there was a significant increase in H_2_-consuming NiFe_1d hydrogenase expression at the community level (**Figure 3A**). Among nine transcriptionally active species expressing NiFe_1d hydrogenases, MGYG000293775, classified as a yet unnamed species of *Duodenibacillus,* dominated in both activity and abundance when methanogenesis was suppressed (**Figure 3B**).

Addressing enteric methane emissions by diverting hydrogen from methanogenesis towards alternative pathways necessitates understanding the competitive interactions among hydrogenotrophic organisms^78^. MGYG000293775 has encoded functional capacity that may provide advantages over other hydrogenotrophs. Notably its flexible capacity for energy generation via anaerobic respiration, coupling hydrogen oxidation to the complete reduction of nitrate or reduction of fumarate to succinate (**Figure 3C and 3E**). Although other hydrogenotrophic genomes encode anaerobic respiratory capabilities, the quinol:fumarate reductase (fdrA/FQR) was unique to MGYG000293775 (**Figure 3C**). Moreover, while nitrate and sulfate have higher free energy potential as terminal electron acceptors^79,80^, their concentrations are typically low in the ruminant diet^20,81^, and likely do not play a major role under the conditions evaluated. Furthermore, the potential for MGYG000293775 to obtain fumarate via DcuABC mediated L-aspartate/L-malate import and succinate export^56^ enables MGYG000293775 to utilize additional rumen metabolites for H_2_/fumarate respiration. These mechanisms, known to confer a growth advantage in other anaerobic systems^56,82^, may provide MGYG000293775 with a competitive advantage over other hydrogenotrophic rumen microbes, and could be potentially exploited in novel approaches for rumen microbiome modulation.

Several strategies have been explored to redirect substrates towards enhanced propionate production in the rumen to reduce methanogenesis^14^. In our study, we uncover mechanisms within MGYG000293775 that enable the efficient conversion of multiple, potentially abundant, rumen metabolites into succinate, a key precursor for propionate synthesis in the rumen^14,55^. Additionally, the mechanism of exporting succinate through the DcuABC-mediated antiporter^56^, and increased ruminal succinate availability, may explain increases in the propionate/acetate ratio following supplementation with *Asparagopsis* sp. previously observed in a semi-continuous *in vitro* rumen model^35^. Although VFAs were not quantified in this study, we noted a reduction in the expression of succinyl-CoA synthetase (*Sus*) and propionyl-CoA carboxylase (*Pcc*) in samples treated with *A. armata* (**Figure 5**). This suppression of propionate biosynthetic genes could potentially be attributed to feedback inhibition, consistent with a scenario where increases in propionate inhibit the enzymes involved in propionate synthesis^83^. Alternatively, a respective decrease in acetate pools would be consistent with our observed upregulation of acetate biosynthetic enzymes (**Figure 5**). Thus, while a promising mechanism for methanogenic substrate redirection has been identified in MGYG000293775, the impact of altered rumen metabolite pools on VFA biosynthetic gene expression requires further investigation to fully understand how metabolic intermediates are redirected in response to perturbing the rumen microbiome with methane-mitigating red seaweed.

While it is well established that excess hydrogen inhibits some of the major cellulolytic rumen bacteria, and therefore could impact the overall degradation of complex carbohydrates in the rumen ecosystem^84^, very little is known about the cellulolytic bacteria that are less sensitive to increased hydrogen and the repertoire of CAZymes they employ under these conditions. Investigating rumen CAZyme expression profiles in our analysis revealed a widespread down-regulation of CAZyme families in *A. armata* treated animals. This downregulation was observed at the CAZyme family level (**Figure 4A**), across most potential CAZyme substrates, and broadly across microbial taxonomic groups expressing these enzymes (**Figures 4B**). Downregulation of CAZyme expression was especially prominent across genomes of the *Clostridia*, whereas significant downregulation of CAZymes in the *Bacteroidia* was limited to those involved in Glucuronan and Pectin metabolism (**Figure 4B and S8**). The observed downregulation of microbial CAZymes did not negatively impact animal feed efficiency (**Table S1**), and notably the single CAZyme family with a significant increase in expression (GH11) has been reported as more highly expressed in feed efficient animals^63^. These findings may indicate that specific and important fibrolytic groups, such as the *Bacteroidia* and *Fibrobacteria* may be generally more resistant to increased concentrations of hydrogen in the rumen, and that those robust bacteria play a more prominent role in biomass degradation under seaweed induced inhibition of methanogenesis. A significant question that remains when developing new methane mitigation strategies is how to further leverage microbiome adaptations that can overcome hydrogen mediated inhibition of biomass-degradation so that feed carbohydrates can be more comprehensively and efficiently utilized by the host animal^85^.

Here we utilized a combination of metagenomics and metatranscriptomics to successfully identify a rumen bacterium, MGYG000293775, with the potential to utilize excess H_2_ for driving the conversion of fumarate to succinate, a key precursor for propionate synthesis, therefore revealing a potential avenue for advanced methane mitigation strategies that redirect methanogenic substrates towards more desired metabolites. Although our study was somewhat limited in sample size, length, animal numbers and sequencing depth and only captured a snapshot of the microbial activity, we were able to obtain new insights into the rumen microbiome and its functional diversity. This highlights the value and importance of expanding these kinds of rumen microbiome studies, across time, locations, breed, etc., and utilizing the latest omics techniques to obtain a more detailed and finely nuanced understanding of the rumen microbiome and especially the functional changes that are triggered by rumen perturbation that aim at a reduced production of enteric methane.

## Materials and Methods

### Experimental design and setup

All procedures were reviewed and approved through the University of California, Davis Institution for Animal Care and Use Committee (IACUC) under protocol number 20398. Details of the animal trial have been described previously by^18^. In brief, three sets of four cows (total n=12) were randomly assigned to one of three treatment groups (Control group: basal diet; Low Dose group: basal diet + 0.5% OM *Asparagopsis armata*; High Dose group: basal diet + 1% OM *A. armata*), then fed the allocated diet for 14 days during which milk production and components, dry matter intake (DMI), body weight (BW), feed conversion efficiency (FCE), CH_4_, carbon dioxide (CO_2_), and hydrogen (H_2_) production were recorded. After the two-week feeding period ended, cows were fed the basal diet for 7 days (washout period), before treatments were randomly reassigned to a different set of cows. This ensured to expose all 12 cows to each of the three treatment groups for a 14-day period. For the work described here, rumen fluid was collected from the Control and High Dose (Treatment) group during the last day of the last period of experiment. An overview of the experimental design is outlined in Supplementary Figure 1.

### Sample collection and preparation

Two hours after feeding, cows were moved to a head gate where rumen fluid was collected using an oral stomach tube technique^86^. An oral-ruminal stomach tube fitted with a perforated brass probe head (Anscitech Co Ltd. Wuhan, China) was inserted through the mouth and into the rumen at an insertion depth of 200 cm to ensure that samples were collected from the central portion of the rumen, providing for less variability between samples^87^. During collection, the first 500 mL of sample was discarded to limit saliva contamination (van Gastelen et al., 2017). Approximately 200 mL of rumen fluid was captured in a 1 gallon prewarmed, insulated canister (Thermos, Schaumburg, IL) and transported immediately to the laboratory. Rumen fluid was strained through four layers of cheesecloth into 100 mL centrifuge tubes, flash frozen in liquid nitrogen and then stored at -80°C to prevent sample degradation.

### DNA/RNA extraction and sequencing

DNA extraction was performed using the FastDNA SPIN Kit for Soil (MP Biomedicals, Solon, OH) with ∼500 mg of sample according to the manufacturer’s protocol. DNA was subsequently purified with a Monarch PCR & DNA Cleanup Kit (New England Biolabs, Ipswich, MA) following the manufacturer’s instructions. Extracted DNA was stored at −20°C until shipped and further processing for DNA shotgun sequencing at the DOE Joint Genome Institute (Walnut Creek, CA).

To obtain RNA, approximately 1 mL of frozen rumen fluid was thawed and homogenized using an 18-gauge needle and syringe., RNA was then extracted and purified using the PureLink RNA Mini Kit (Invitrogen, San Diego, CA). After purification, the samples were then treated with DNA-Free Kit DNase treatment and removal (Invitrogen, Waltham, Massachusetts, USA) at room temperature for 15 minutes. RNA was quantified using a Bioanalyzer (Agilent, Santa Clara, CA) and RNA quality was evaluated using a sodium hypochlorite (6%) agarose gel as described previously (Aranda et al., 2012). All RNA samples were equally split in half then subjected to two different ribosomal RNA removal techniques [i.e., Ribo-Zero(TM) rRNA Removal Kit (Illumina, San Diego, CA, USA) and QIAseq FastSelect rRNA Removal Kit (Qiagen, Germantown, MD]. Both RNA sample quantities were determined using Qubit 3.0 Fluorometer (Invitrogen, Waltham, Massachusetts, USA)). JGI, RiboZero: rRNA was removed from 10 ng of total RNA using Ribo-Zero(TM) rRNA Removal Kit (Illumina, San Diego, CA, USA). Stranded cDNA libraries were generated using the Illumina Truseq Stranded mRNA Library Prep kit. The rRNA depleted RNA was fragmented and reversed transcribed using random hexamers and SSII (Invitrogen, Waltham, MA, USA) followed by second strand synthesis. The fragmented cDNA was treated with end-pair, A-tailing, adapter ligation, and 8 cycles of PCR. The prepared libraries were quantified using KAPA Biosystems’ next-generation sequencing library qPCR kit and run on a Roche LightCycler 480 real-time PCR instrument. Sequencing of the flowcell was performed on the Illumina NovaSeq sequencer using NovaSeq XP V1 reagent kits, S4 flowcell, following a 2×151 indexed run recipe. JGI, FastSelect: The prepared libraries were quantified using KAPA Biosystems’ next-generation sequencing library qPCR kit and run on a Roche LightCycler 480 real-time PCR instrument. Sequencing of the flowcell was performed on the Illumina NovaSeq sequencer using NovaSeq XP V1 reagent kits, S4 flowcell, following a 2×151 indexed run recipe.

### Recovery of rumen metagenome-assembled genomes (RuMAGs)

The two metagenomes were subject to quality control with FastQC (http://www.bioinformatics.babraham.ac.uk/projects/fastqc). The adaptors and PhiX contaminants were detected and trimmed by bbMap^88^ followed by quality trimming with sickle^89^ with default parameters. The high-quality reads were assembled with IDBA (--mink 30 --maxk 150 --step 10)^90^. The contigs longer than 5000bp were extended with COBRA^91^ by the identification of overlapped kmers at both 5’ and 3’ terminus among all the contigs and consensus coverages across the joint contigs which was estimated by mapping the reads against the contigs with bbMap. The contig coverage was calculated using the jgi_summarize_bam_contig_depths script from the MetaBAT2 package^92^.

In order to recover the MAGs from the contigs, cross-mapping of the two sets of metagenomic reads against the two assemblies (contigs >=2.5kb), was performed with SNAP^93^ and the contig coverage was calculated by jgi_summarize_bam_contig_depths. The coverage file and assemblies were used to bin assembled contigs into genome bins (MAGs) using concoct^94^, maxbin2^95^, metabinner^96^, metabat2^92^ and vamb^97^. To remove the redundancy of MAGs and pick up the best-quality MAGs among different binners, DASTool^98^ was deployed (--score_threshold 0.3), which resulted in 226 non-dereplicated MAGs across the two samples analyzed using metagenomics.

### Construction of non-redundant rumen prokaryotic genome database

We identified and integrated additional microbial genomes from rumen samples from publicly available databases and integrated them with the MAGs recovered in this study. This included 457 isolate genomes from the Huntage1000 collection^40^, 4941 MAGs from^41^, and 5578 genomes from the Mignify database^42^. Genomes from each set were individually de-replicated at the species level with dRep^99^ (--S_algorithm ANImf -sa 0.95 -nc 0.10 -comp 60 -con 10). This step resulted in 345, 2177, 2729 and 72 non-redundant species-level genome representatives (ANI ≥ 95 %) from the Hungate1000, RUGs, and Mignify collections and MAGs from this study. Finally, the species-level representative genomes from each database were integrated and de-replicated at the species-level (ANI ≥ 95 %) with dRep a second time to generate a final comprehensive database of 3,180 non-redundant species-level representative genomes. The taxonomic classification of the genomes was conducted with GTDB-TK^100^. To visualize the phylogenetic relationships among the genomes, the bacterial and archaeal trees were constructed with GToTree^101^ by designating the bacterial and archaeal single-copy marker gene sets separately and trees were visualized with iTol^102^.

### Functional annotations of genomes

We employed several different approaches for genome functional annotation. KofamScan^103^ (KofamKOALA, -T 0.75) and eggNOG-mapper v2^104^ were used for general functional annotation of genomes. If multiple HMM profiles from KofamScan matched a gene, only the highest scoring hit was retained. Carbohydrate-active enzymes were predicted by using dbCAN3^105^.

Custom hidden markov models (HMMs) were generated for hydrogenase classification. Briefly, the reference sequences from each group of hydrogenase within HydDB^48^ were retrieved and multiple alignment was performed separately for each group of protein sequences using Clustalo^106^. This included major hydrogenase groups FeFe-1, FeFe-2, FeFe-3, FeFe-4, NiFe-A, NiFe-B and NiFe-C. HMM models were constructed from each protein sequence alignment using the hmmbuild function of the HMMER package^107^. Reference sequences from all groups were scored against the resulting HMM models with hmmsearch, and HMM score thresholds for identifying group membership were set as 75% of the score from the lowest scoring reference sequence known to belong to the respective group. Both an E-value <1e-15 and score threshold were used to filter the results when searching genomes against the hydrogenase HMM models with hmmsearch. To better annotate the predicted hydrogenases into well-defined subgroups (e.g., NiFe_1a, NiFe_1b, NiFe_1c), manual phylogenomic annotation was deployed. The predicted hydrogenases and reference hydrogenases from the same group were integrated and aligned with Clustalo. Phylogenetic tree was constructed by Fasttree^108^. The subgroups for predicted hydrogenases were defined by their closest reference hydrogenases. This well classified the subgroups for NiFe-1, NiFe-2, NiFe-3, NiFe-4 and FeFe-C. There’s a set of NiFe-4 hydrogenase closest to NiFe-4e but no references were found within the clade itself, so we labeled them as ‘NiFe-4e-potential’ (**Figure S9**). For FeFe-A hydrogenases, FeFe-A2 and FeFe-A4 could be classified on the tree but FeFe-A1 and A3 could not be well separated. FeFe-A3 hydrogenases have been reported to be coupled with NADH-dependent oxidoreductases^47^, so the flanking genes of those unclassified FeFe-hydrogenases were extracted and the hydrogenases accompanied by NADH-dependent dehydrogenases or reductases were assigned to FeFe-A3 with the remainder assigned to FeFe-A1. For Fe-only hydrogenase, there are only 25 reference sequences in HyDB so we did not intend to build an HMM model for it. Rather we used KO annotation to identify Fe-only hydrogenases in our genomes since the 25 reference sequences all belong to KO entry K13942 in the KEGG database. The list of potential reductases receiving the electrons from hydrogen can be found in **Table S9**.

### Identification of methanogenesis and VFA synthesis pathways

The enzymes involved in methanogenesis pathways from methanol, acetate, methylamine and CO_2_, acetate and propionate biosynthetic pathways were identified based on literature search and the corresponding KOs for these reactions were retrieved from the KEGG database^109^. For acetate and propionate biosynthesis pathways, we retrieved the corresponding KOs from the literature^110–113^. For butyrate biosynthesis, we leveraged the hmm model introduced here^67^ to identify the corresponding genes. For each reaction, if one of the subunits is present in the genome, we assume this genome is capable of catalyzing this reaction. For the presence of a complete biosynthetic pathway, we only allow 2 missing reactions maximum. The rule of co-localization of butyrate biosynthesis-related genes was not applied here because of the fragmented scaffolds in MAGs. For a detailed description of the KO list involved in methanogenesis and VFA synthesis and criteria used to define the presence of a specific pathway, please refer to **Table S10**.

### Calculation of the genome coverage

To calculate the genome coverage, we mapped the reads from metagenomic samples against our comprehensive genome database using bowtie2^114^. Subsequently the sample specific coverage of individual genomes was calculated using CoverM (--min-read-percent-identity 0.95 --min-read-aligned-length 50). To filter out the spuriously detected genomes due to ambiguous alignment against conserved regions among different genomes, we calculated the expected breadth of a genome as a function of its coverage with the following formula^115^:

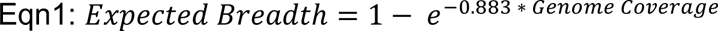

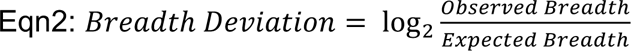

The coverage of any genome from a single sample where < 100 reads were mapped to the genome or where the value Breadth Deviation < -1 was changed to zero in our analysis and was considered not conclusively identified as present in the sample.

### Metatranscriptomic data analysis

The metatranscriptomic reads were subject to quality trimming in the same way as metagenomic reads as described above. rRNA reads were detected and depleted with sortmerna^116^. The reads from the same sample while being generated from libraries prepared with different rRNA depletion methods were combined given the fact that they should capture highly consistent transcripts (**Figure S10**). To profile the transcript expression level, reads were mapped against the predicted ORFs in each genome with bowtie2^114^. Only mapped reads with a MAPQ score ≥35 were retained. Expression levels were calculated by CoverM (--min-read-percent-identity 0.97 –min-covered-fraction 0.95, https://github.com/wwood/CoverM) and only genes covered by at least 5 reads were retained. Differential expression was conducted with DESeq2^117^. The variance stabilizing transformed expression profile by DESeq2 was used to conduct principal component analysis using the prcomp function in R. To assess the differences in overall transcriptome profiles between control and treatment samples PERMANOVA was conducted based on *Bray Curtis* dissimilarity of the transformed expression profile with VEGAN package^118^.

## Data Availability

Raw metagenomic and transcriptomic data used in this study are available from the JGI Genome Portal (https://genome.jgi.doe.gov/portal/). JGI Project ID numbers for each sample can be found in Supplementary Table S2. Non-redundant species representative genomic bins generated in this study will be made available under a pending NCBI bioproject. All data will be made available upon request.

## Acknowledgements

DNA and RNA library construction and subsequent sequencing was performed under proposal: 10.46936/10.25585/60001243 by the U.S. Department of Energy Joint Genome Institute (https://ror.org/04xm1d337), a DOE Office of Science User Facility, is supported by the Office of Science of the U.S. Department of Energy operated under Contract No. DE-AC02-05CH11231. Funding for this project was also provided in part by the Shurl and Kay Curci Foundation. This work was also supported in part by Lyda Hill Philanthropies, Acton Family Giving, the Valhalla Foundation, Hastings/Quillin Fund - an advised fund of the Silicon Valley Community Foundation, the CH Foundation, Laura and Gary Lauder and Family, the Sea Grape Foundation, the Emerson Collective, Mike Schroepfer and Erin Hoffman Family Fund - an advised fund of Silicon Valley Community Foundation, the Anne Wojcicki Foundation through The Audacious Project at the Innovative Genomics Institute.

## Author Contributions

Conceived, designed, and performed the experiments: BR, EK, MH. Generated data: NS, EE; Analyzed the data: PZ, BR, PR, SD, MH. Wrote major parts of the manuscript: PZ, SD, MH. Contributed to the manuscript: PR, EK. Reviewed and approved manuscript: all authors.

## Corresponding Author

All correspondence should be addressed to Matthias Hess

