## Supplementary Figures for "Red seaweed supplementation suppresses methanogenesis in the rumen, revealing potentially advantageous traits among hydrogenotrophic bacteria"

### Red seaweed supplementation suppresses methanogenesis in the rumen, revealing potential competitive strategies among hydrogenotrophic bacteria

Pengfan Zhang[1], Breanna Roque[2,3], Pedro Romero[2], Nicole Shapiro[4], Emiley Eloe-Fadrosh[4], Ermias Kebreab[2], Spencer Diamond[1], Matthias Hess[2,*]

[1]Innovative Genomics Institute, University of California, Berkeley, CA, USA.
[2]Department of Animal Science, University of California, Davis, CA, USA.
[3]FutureFeed, Garbutt, Queensland, Australia
[4]DOE Joint Genome Institute, Berkeley, CA, USA.

***Corresponding author:**

Matthias Hess
University of California, Davis
Department of Animal Science
2251 Meyer Hall
Davis, CA 95616, USA

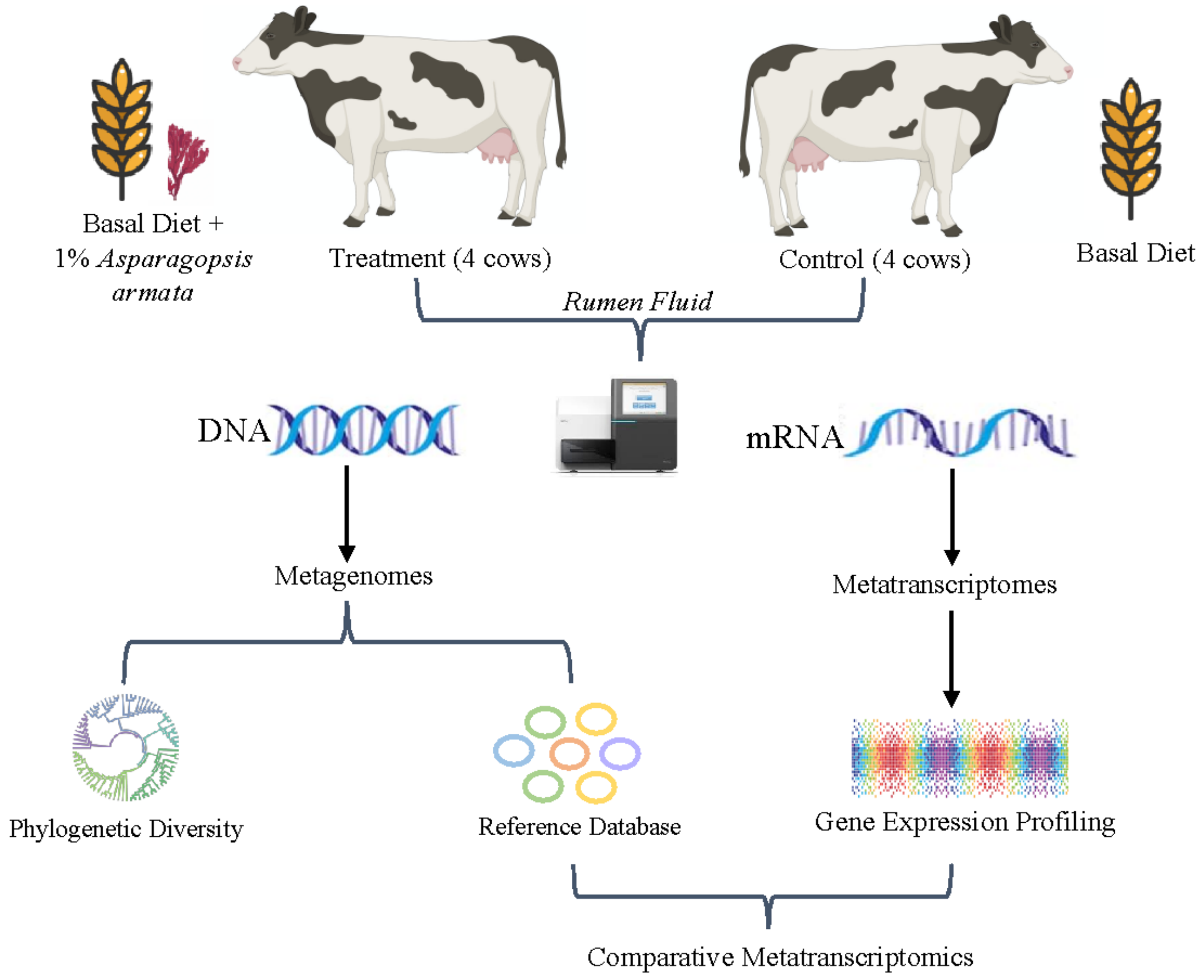


**Supplementary Figure 1: Experimental design of this study.** Cows were fed with a basal diet (control; n = 4 animals) or with a basal diet supplemented with 1% *Asparagopsis armata* (treatment; n = 4 animals). The rumen fluid of the animals was collected after 14 days and was subsequently subjected to metagenomic and metatranscriptomic sequencing.

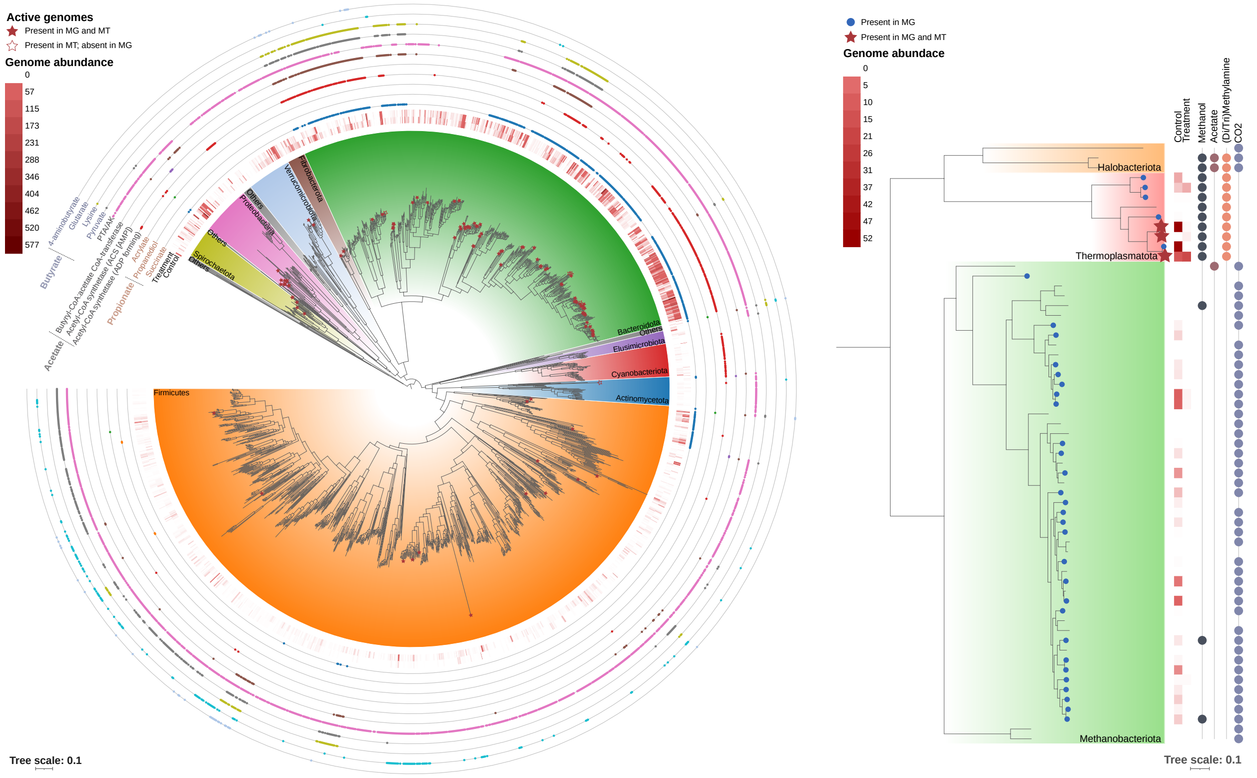


**Supplementary Figure 2: Phylogenetic tree of 3,180 rumen bacteria and archaea. A**) Phylogenetic tree of 3,075 bacterial genomes in the rumen specific database created for this study at species level resolution. Tree was constructed using a concatenated set of bacterial specific marker genes, and rooted at the midpoint. Some genomes were omitted from the tree due to insufficient numbers of marker genes. Colored ranges over tree branches indicate phylum level taxonomy. Phyla that had fewer than 20 species were consolidated in “Others”. Solid stars indicate genomes for which were detected in metagenomic samples and ≥ 20% of genes were detected in metatranscriptome samples. Non-solid stars indicate genomes that recruited reads from metatranscriptome samples, but were not detected in metagenome samples. The heatmap shows normalized genome abundance within the metagenome samples as indicated in the inner two rings (for Control and Treatment samples). The outer three rings indicate the presence of VFA (e.g. propionate, acetate and butyrate) biosynthetic pathways in genomes by the presence of colored circles. **B**) Phylogenetic tree of all 61 archeal genomes from the methanogenic orders *Halobacteriota*, *Thermoplasmatota* and *Methanobacteriota* in the rumen specific database created for this study at species level resolution. Tree was constructed using a concatenated set of archaeal specific marker genes, and rooted at the midpoint. Solid stars indicate genomes for which were detected in metagenomic samples and ≥ 20% of genes were detected in metatranscriptome samples. The blue dots indicate the genomes were only detected in metagenomic samples. The heatmap shows normalized genome abundance within the metagenomes as indicated in the inner two rings (for Control and Treatment samples). Colored points below the heatmap indicate the presence in genomes of methanogenic metabolic pathways that can utilize different starting substrates.

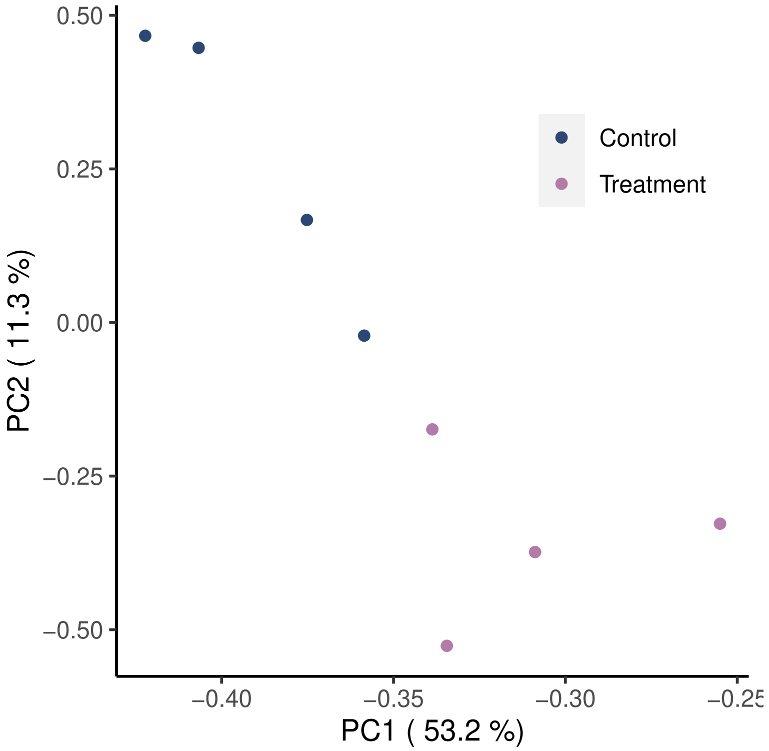


**Supplementary Figure 3: Compositional differences in transcriptome profiles of control and treated animals.** Principal component analysis (PCA) was performed based on the variance-stabilizing-transformed transcript counts (DESeq2) of KEGG Orthology Groups (KOs) in each sample. Significant differences in KO abundance composition between samples based on treatment was evaluated using permutational analysis of variance (PERMANOVA) based on the *Bray Curtis* dissimilarity of the transformed transcript counts (p-value = 0.034).

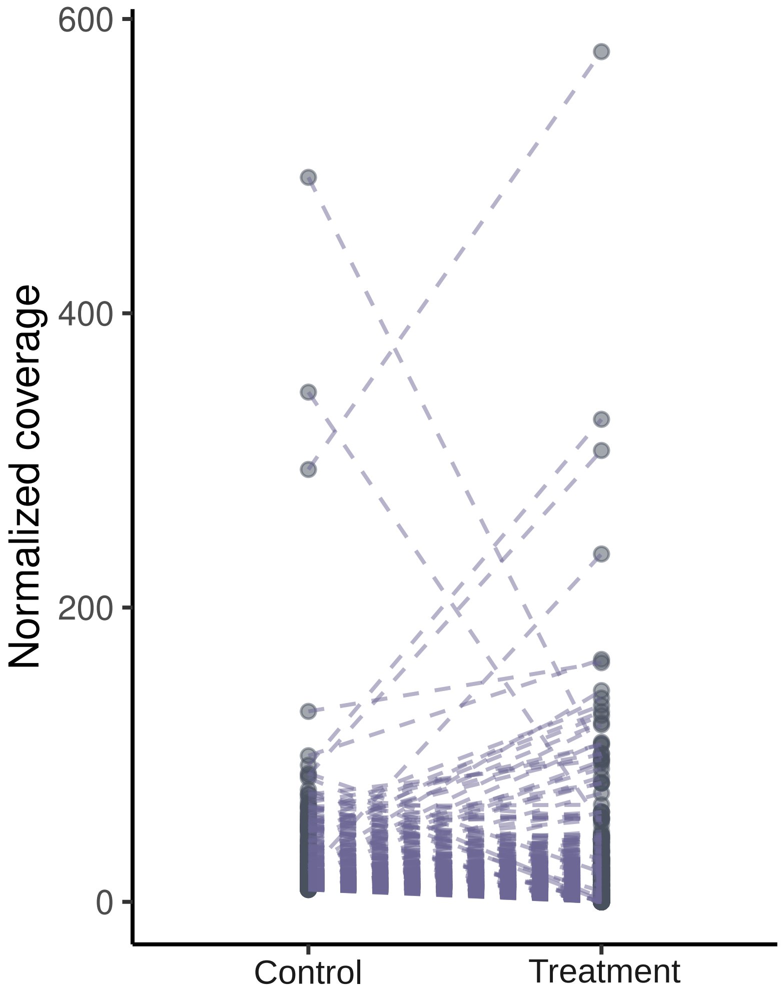


**Supplementary Figure 4: Abundance shifts for individual genomes in the rumen microbiome after A. *armata* treatment.** The top 250 most abundant genomes and their corresponding sequencing depth-normalized coverage are shown. The dashed line connects the coverages of the same microbial genome from the rumen of control and treated animals.

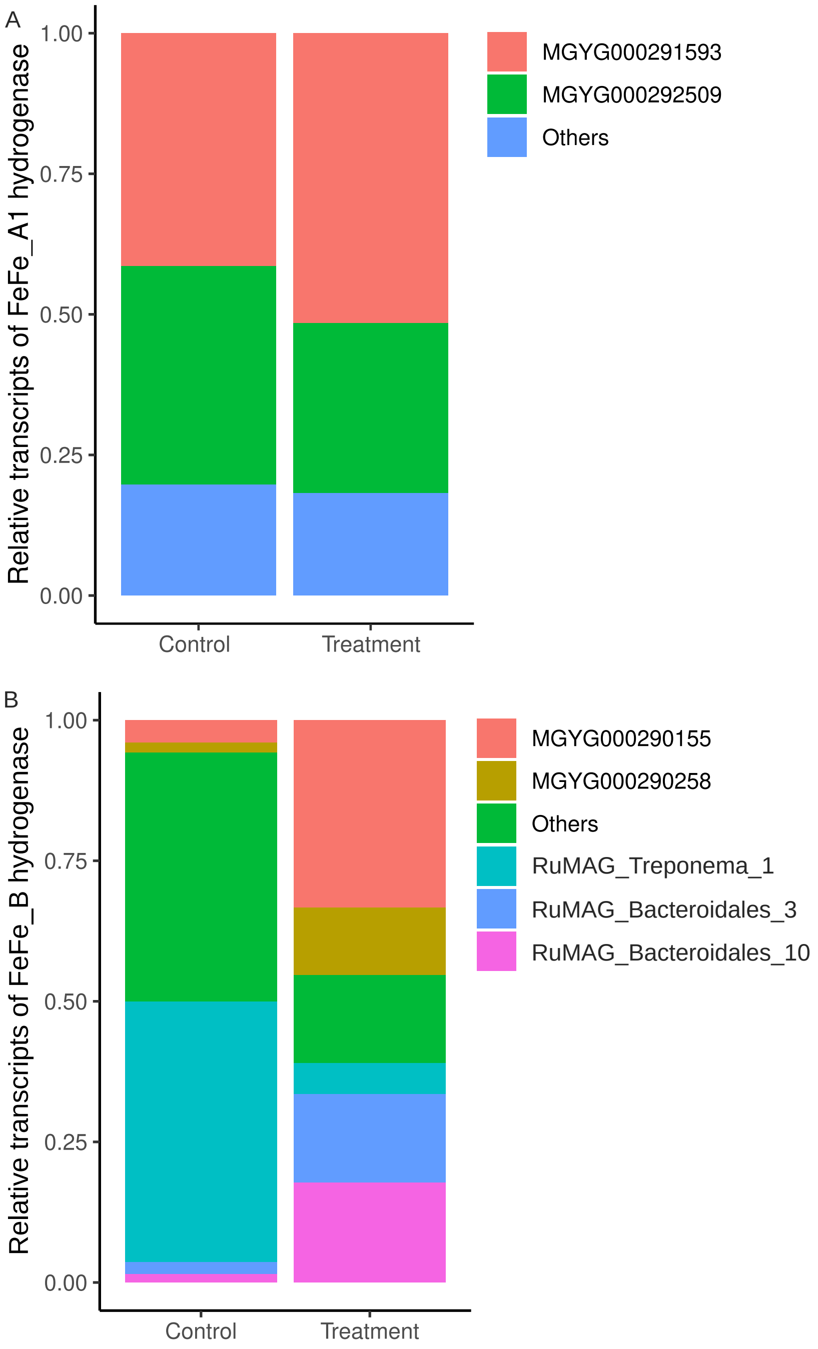


**Supplementary Figure 5: Species-level contributions to transcripts of hydrogen evolving FeFe-hydrogenases. A)** The fractional contribution of transcripts assigned to FeFe_A1 hydrogenases in treatment and control samples coming from individual species. Data is aggregated across the 4 treatment and control samples, respectively. **B)** The fractional contribution of transcripts assigned to FeFe_B hydrogenases in treatment and control samples coming from individual species. Data is aggregated across the 4 treatment and control samples, respectively. Colors indicate the genome from where transcripts were derived. Only the most active contributing genomes are displayed and the other genomes are consolidated in “Others”.

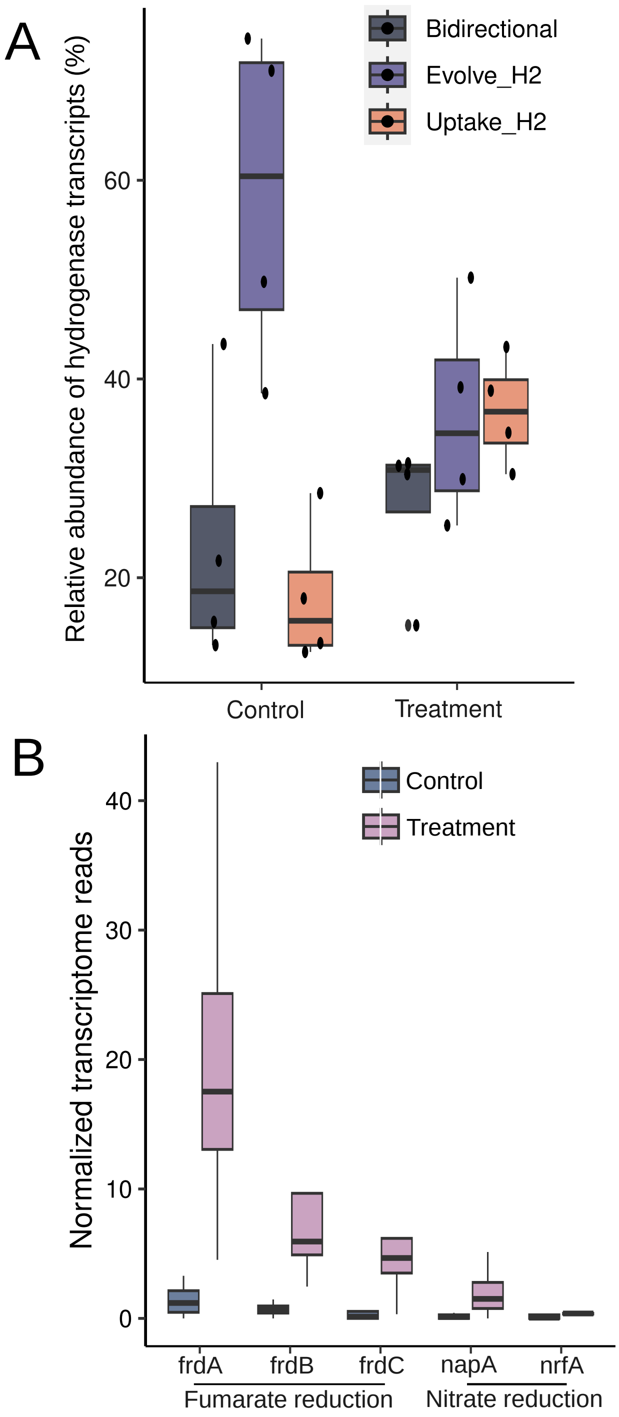


**Supplementary Figure 6: Hydrogenase expression compositional profile of animals and gene expression changes in MGYG000293775 in response to A. armata treatment. A**) The y-axis shows the compositional abundance of transcripts derived from three hydrogenase categories: hydrogen-consuming, hydrogen-uptakem and bidirectional hydrogenases in the rumen microbiome collected from either control or treated animals. Data is aggregated across the 4 treatment and control samples, respectively. **B**) The transcript abundance of fumarate quinol (FQR) and nitrate reductases identified in the genome MGYG000293775. frdA-C: fumarate quinol reductase A-C; napA: nitrate reductase; nrfA: nitrite reductase.

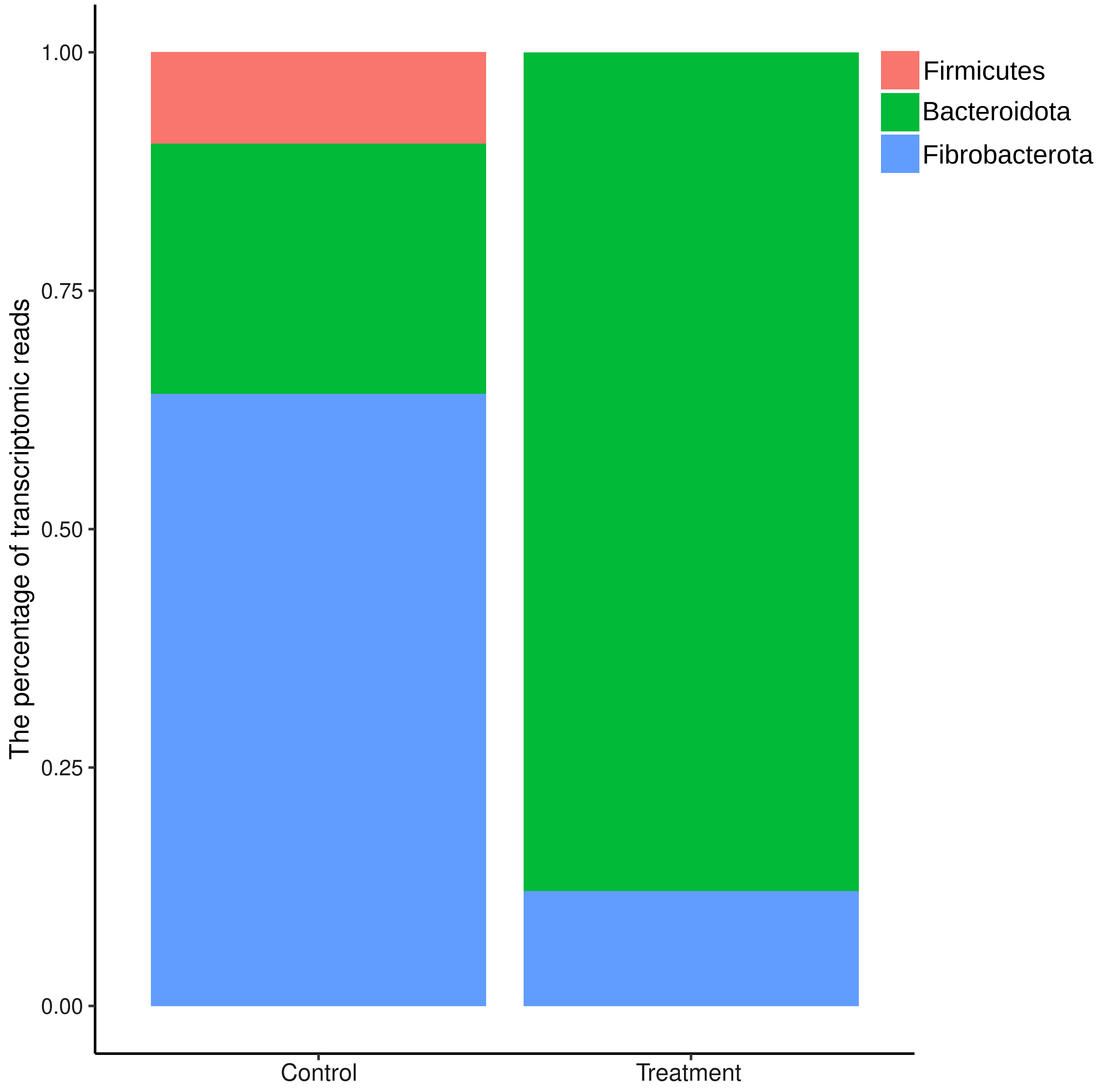


**Supplementary Figure 7: Relative contributions of bacterial phyla to GH11 transcripts.** The y-axis shows the relative GH11 transcript reads contributed by each phylum in the rumen microbiome from either control or treated animals.

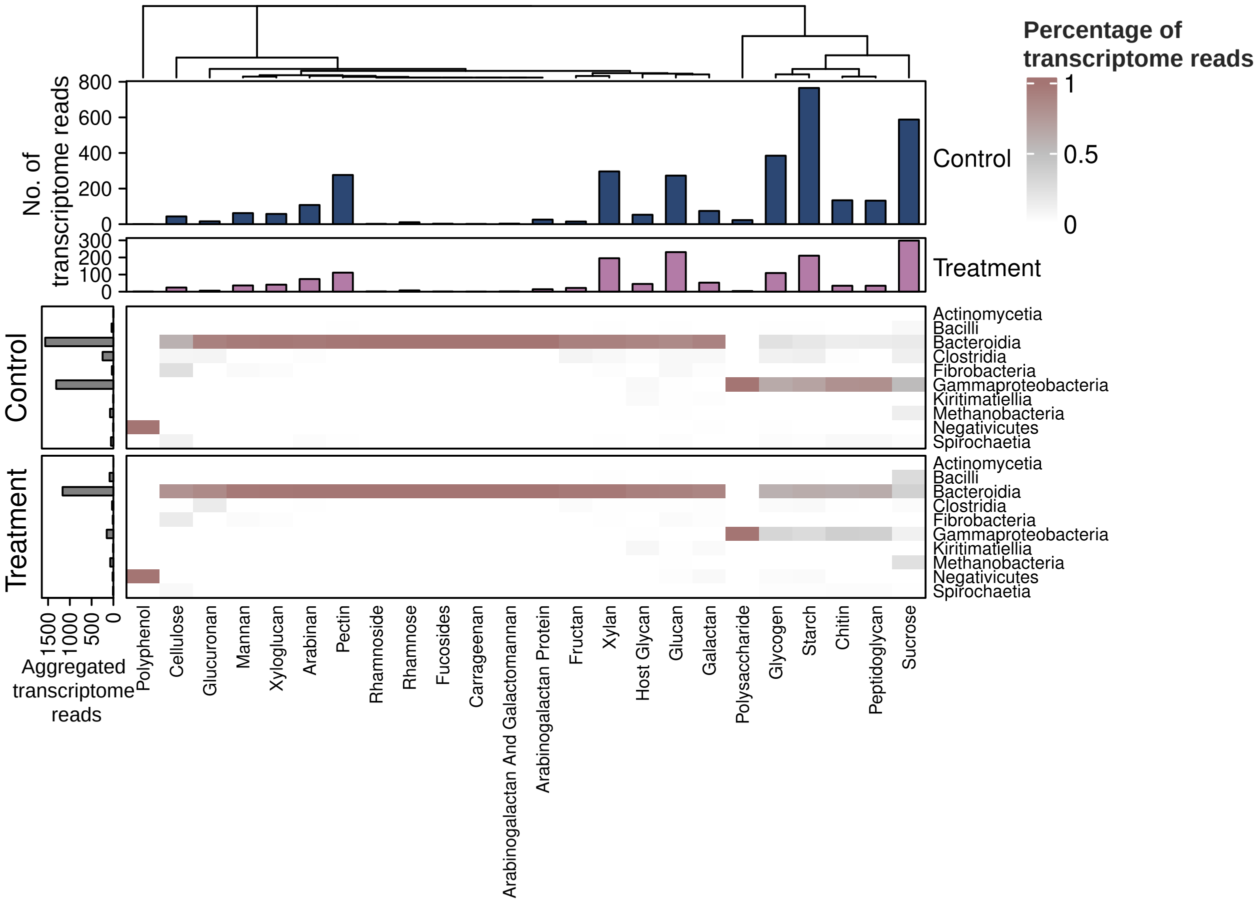


**Supplementary Figure 8: Taxonomic contribution to the degradation of diverse carbohydrates.** The heatmap shows the relative contribution of each bacterial class to the transcript reads of the CAZymes characterized by their corresponding carbohydrate targets. The left bar plot shows the aggregated CAZyme transcript reads (sequence depths normalized) in each class. The two top bar plots indicate the aggregated CAZyme transcript reads (sequence depths normalized) by carbohydrate targets in the rumen microbiome from either control or treated animals.

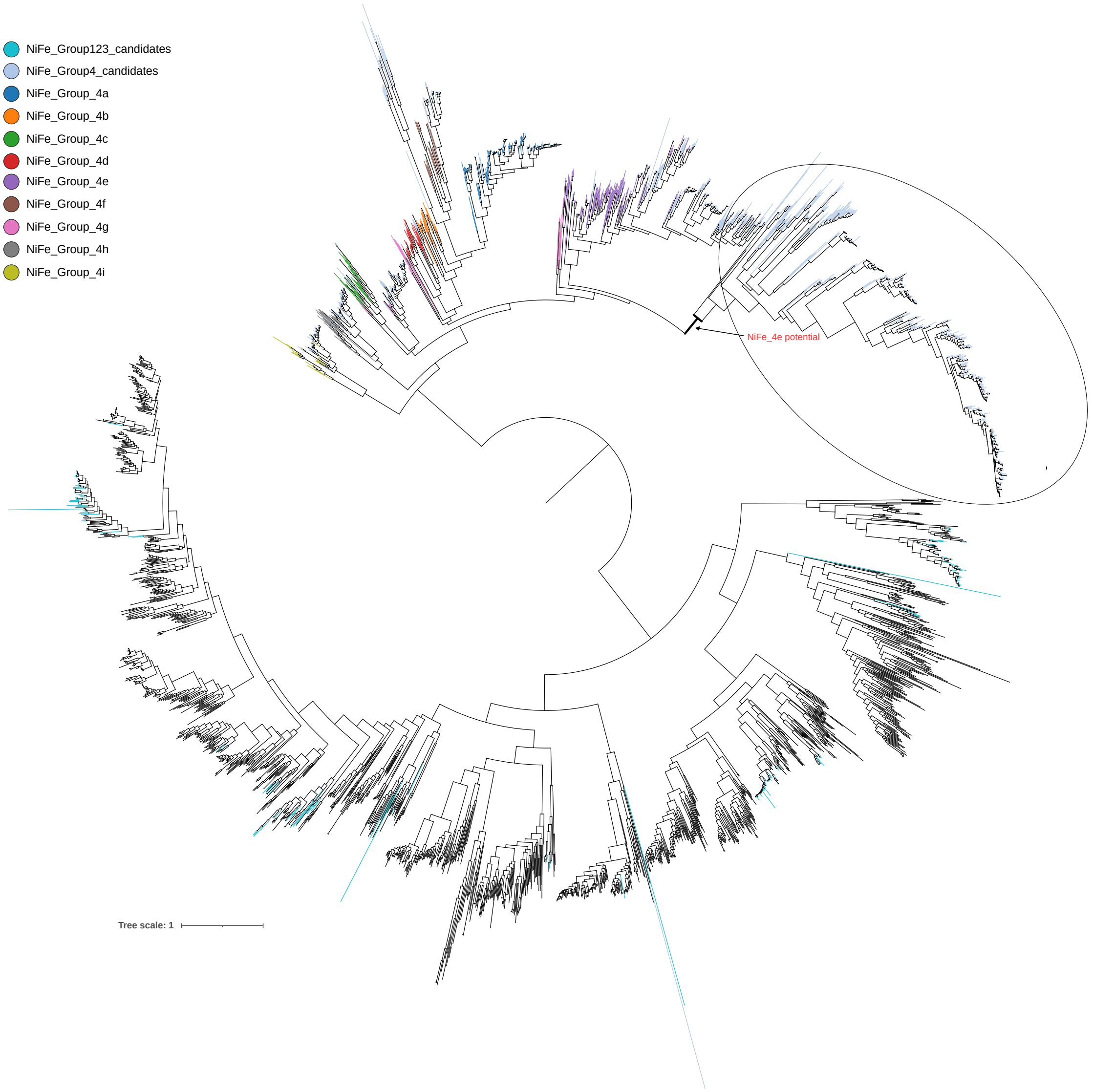


**Supplementary Figure 9: The phylogenetic tree of hydrogenases.** The NiFe-hydrogenases identified in the rumen specific genome database and in the HyDB reference database were aligned by Clustalo and the tree was constructed with Fasttree. Only the reference hydrogenases within the NiFe_4 subgroup and novel hydrogenases identified in the rumen specific genome database are colored. The potential NiFe_4e hydrogenases are highlighted within the ellipse.

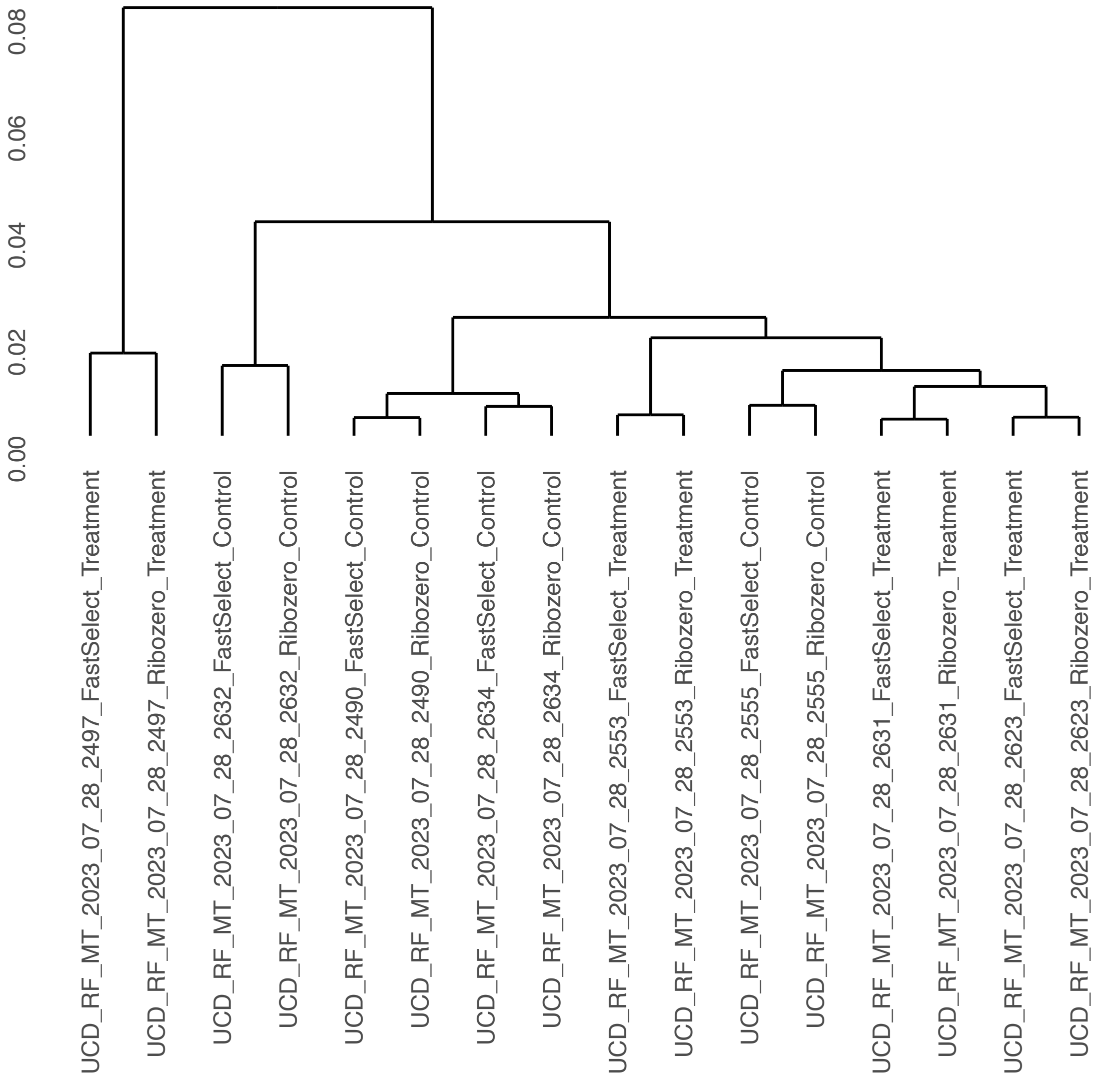


**Supplementary Figure 10: Consistent transcriptional profiles derived from two different RNA depletion strategies.** The gene-level transcriptional profiles generated from two different RNA depletion methods (FastSelect and Ribozero) were clustered with `hclust` function in R. The samples from the same animal are consistently clustered together and are independent of RNA depletion strategies employed.
